# SHANK3–CAMSAP2 interaction links synapses to dendritic microtubule organisation in PV neurons

**DOI:** 10.64898/2026.02.10.705051

**Authors:** Daniela Hacker, Suho Lee, Michael Hecht-Bucher, Sangmun Lee, Caspar Lamborelle, Alois Nathanael Doil, Tomas Fanutza, Arie M. Brueckner, Taesun Yoo, Sacnicte Ramirez-Rios, Cezary Smaczniak, Kerstin Kaufmann, Dick H. W. Dekkers, Jeroen A. A. Demmers, Marie-Jo Moutin, Eunjoon Kim, Marina Mikhaylova

## Abstract

The microtubule cytoskeleton plays an essential role in establishing and maintaining neuronal polarity. In neurons, microtubules are initially generated at the centrosome which gradually loses this role in development. In mature neurons, microtubules are stabilised by the microtubule minus end-binding protein CAMSAP2. How and where microtubule minus ends are anchored within dendrites of mammalian neurons remains an open question. Here, we show that microtubules are directly attached to the postsynaptic density of excitatory synapses through an interaction of CAMSAP2 with the scaffolding protein SHANK3. This process is particularly relevant in parvalbumin-positive GABAergic neurons, in which excitatory synapses are predominantly located directly on the dendritic shaft in direct proximity to microtubules. Notably, this association is strongly enhanced by the Autism Spectrum Disorder (ASD)-associated SHANK3_L68P_ mutation, leading to an increase of synaptic levels of CAMSAP2. This promotes an increased microtubule association with the synapse and alters microtubule dynamics. Downregulation of CAMSAP2 in SHANK3L68P parvalbumin neurons tips the balance between site-specific microtubule stabilisation and dynamics, reshaping dendritic architecture, connecting SHANK3-CAMSAP2 dependent microtubule regulation to synaptic ASD pathology.

**Teaser:** Interaction between synapses and microtubules is enhanced in parvalbumin neurons bearing the ASD-associated SHANK3 L68P mutation.

## Introduction

Long-lived postmitotic neurons rely on a complex microtubule (MT) cytoskeleton network that is integral for shaping its vast dendritic arbour and axonal processes, determining neuronal polarity and mediating polarised long-distance cargo trafficking. Dynamic instability of MTs, which cycle between growth and shrinkage (*1*), mediates neuronal growth, differentiation, and dendritic branching as well as a neuron’s ability to adapt to changes (*2*). MT dynamics are influenced by MT-associated proteins binding to the MT plus or minus end or the MT lattice (*3*). Both dynamics and stability of MTs can be regulated by MT associated proteins that bind to their minus end, such as Calmodulin-regulated spectrin associated protein 2 (CAMSAP2) (*4*). A common read-out of MT stability is the presence of post-translational modifications (PTMs): stable MTs are typically acetylated and detyrosinated, while dynamic MTs are tyrosinated (*5*). In excitatory neurons, peripheral MTs near the plasma membrane of neurites are considered dynamic, whereas MTs in the centre of the dendrite are regarded as stable (*6*, *7*). Since MTs are polarised structures and molecular motors move either towards their plus or the minus end, MT orientation plays a vital role in establishing neuronal polarity and in directing transport processes (*7*, *8*).

Our knowledge of MT polarity in neurons of the mammalian brain primarily comes from cell culture experiments, in which excitatory hippocampal or cortical neurons have been studied (*9–12*). In the axon, MTs are organised with their plus ends pointing to the cell periphery (*13*) and the polarity is thought to be determined through MT nucleation occurring at the somatic cis-Golgi (*14*) and subsequent stabilisation of nucleated MTs at the trans-Golgi (*15*). In dendrites of excitatory neurons, MTs exhibit mixed orientation (*7*, *12*, *13*). In developing neurons, MT minus ends are often anchored at the γ-tubulin ring complex of the centrosome that provides MT nucleation from the soma (*16*). During neuronal development, the centrosome’s function is lost and instead non-centrosomal MTs emerge (*17*). In *Drosophila,* membranous organelles such as Golgi outposts, can be found at dendritic branching points have been identified as nucleation sites (*18*, *19*), thought to maintain MT orientation. In dendrites of *Drosophila* neurons, microtubules are however generally oriented with the minus end out (*20*) which is a stark contrast to the mixed polarity exhibited by mammalian microtubules found in dendrites. It remains an open question where dendritic MTs are nucleated and anchored in mammalian brain neurons, because Golgi outposts are largely present in the proximal region and at branch points of the apical dendrite (*21*), but do not appear to be essential for MT nucleation.

Although MTs are rarely found in dendritic spines which contain most excitatory synapses, spine stability and growth are often associated with transient MT entry of the dynamic plus end (*22–24*). CAMSAP2 frequently associates with minus ends of MTs across the dendritic arbour of excitatory neurons (*25*), however minus ends of MTs have not been found in dendritic spines to date.

While most postsynaptic sites of excitatory synapses are located on dendritic spines, aspiny GABAergic neurons receive most of their excitatory postsynaptic input directly on the dendritic shaft in so-called “shaft synapses” (*26*). In the hippocampus, the majority of aspiny neurons are parvalbumin (PV) positive interneurons. They are comprised of basket and chandelier cells, known to be fast-spiking and can be subdivided into two distinct morphologies in different regions of the hippocampus (*27*): PV neurons in the dentate gyrus display dendrites with numerous dendritic spines (*28*); PV neurons within the CA1 region generally exhibit aspiny dendrites that receive excitatory input directly on the dendritic shaft (*28*). Their dendritic arbours are less branched than those of excitatory neurons (*29*).

A central part of the excitatory postsynapse is the membrane-associated postsynaptic density (PSD). It serves as an anchoring platform for membrane receptors (*30*, *31*) such as glutamate receptors, ion channels and adhesion molecules. Excitatory PSDs are organised by scaffolding proteins such as PSD95 and SH3 And Multiple Ankyrin Repeat Domains (SHANK) protein family members (*32*). SHANK proteins link the PSD to F-actin which is involved in shaping spine morphology and influencing their dynamics (*32*, *33*). For the SHANK protein family consisting of three members SHANK1-3, a multitude of alterations including missense mutations are described that are associated with Autism Spectrum Disorder (ASD) or Phelan McDermid syndrome (*34–36*). Some known ASD-related SHANK3 mutations can alter spine morphology by influencing the F-actin cytoskeleton (*37*). Synaptic morphology can thus be influenced by the stability of synaptic scaffolding proteins. In some cases, the protein carrying an ASD-associated point mutation can be less stable compared to SHANK3_WT_ (*38*, *39*), highlighting the effect that single point mutations can exert on protein stability.

The localisation of synapses in PV neurons places excitatory postsynapses in close physical proximity to the dendritic MT cytoskeleton, enabling a link between the MT network and the postsynaptic density. Presently, the MT organisation of PV neurons remains unknown, leaving the potential interaction between MTs and the postsynapse unexplored.

In this study, we investigate the MT organisation of PV neurons and found that MTs are anchored to excitatory PSDs. We provide mechanistic insights on how the association between the MT minus end binding and stabilising protein CAMSAP2 and the scaffolding protein SHANK3 impacts the dendritic cytoskeleton and find that the ASD-associated L68P mutation of SHANK3 results in a gain of function such as an increased association of MTs with the PSD. This impacts dendritic complexity of PV neurons, possibly contributing to the ASD-pathology.

## Results

### The PSD of excitatory synapses anchors microtubules in PV-positive GABAergic interneurons

The organisation of the MT cytoskeleton influences synaptic transmission (*40*, *41*), through its pivotal role in the transport of cargo to and from the synapse (*42*). While most interactions of MTs with the synapse focus on the presynapse, or on postsynapses located on dendritic spines with the MT plus end (*23*, *40*, *41*, *43*, *44*), much less is known about the role of MTs in neurons that predominantly display excitatory synapses directly on the dendritic shaft in close proximity to the MT cytoskeleton. Therefore, we asked what spatial relationship exists between the cytoskeleton and the postsynapse at aspiny PV neurons. We decided to use dissociated cultures which are commonly chosen as an optically accessible model to investigate the cytoskeleton and synaptic organisation at the micro- and nanoscopic level (*45*). This model system is however mainly used to investigate excitatory neurons which make up 89 to 94.2% of the rodent hippocampus (*46*, *47*).

To assess whether dissociated primary hippocampal cultures are suitable to investigate the cell biology of GABAergic neurons, we measured the share of various GABAergic subtypes characterised through the expression of marker proteins such as PV, calbindin or calretinin. In mature cultures (∼day *in vitro* (DIV)15), we found that 8% of microtubule associated protein 2 (MAP2)-positive neurons were also positive for calbindin, 18% for calretinin and 8% for PV (***Fig***1A-B, ***Fig* EV**1A-B). This is comparable to interneuron subtype abundance studies in mouse brains which estimate that around 3.2% to 6% of neurons are PV+ (*46*, *48*). In rat hippocampus, 5.8 – 11% of neurons are estimated to be GABAergic (*46*, *47*) which is comparable to observations we made in a previous study (*49*). We thus concluded that dissociated hippocampal cultures are a suitable model system for our study.

**Figure 1:**
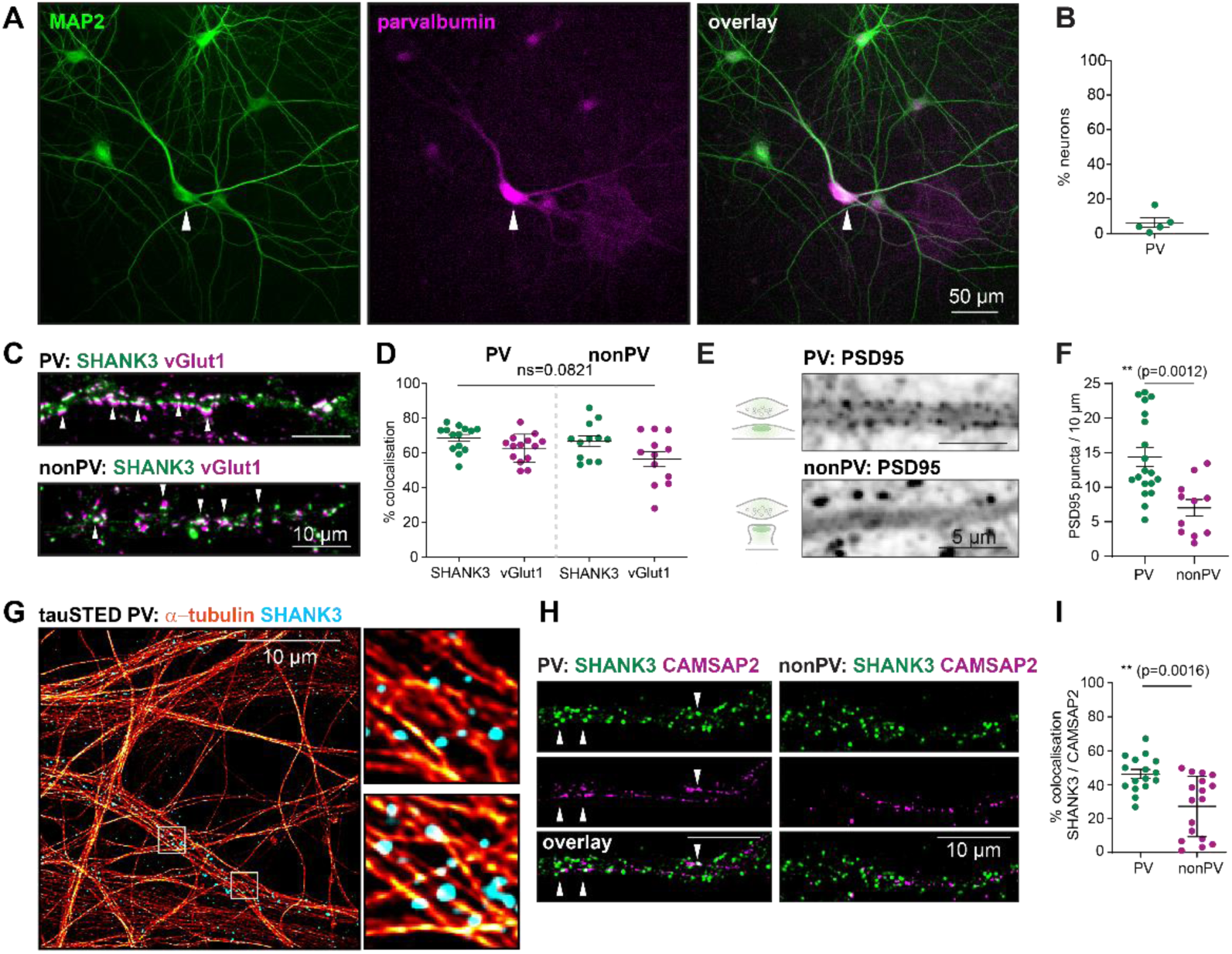
The microtubule cytoskeleton is found at excitatory postsynaptic densities of parvalbumin neurons. **A**: Dissociated hippocampal neuron culture (DIV15) immunostained for the neuronal marker MAP2 (green) and parvalbumin (PV). Arrow indicates PV neuron. Scale bar, 50 μm. **B**: Quantification of 247 fields of view (fov, 655×655 mm^2^; N = 5 independent cultures) stained as in (**A**) to determine the share of PV neurons compared to the total number of neurons. **C**: Dissociated hippocampal neuron culture (DIV15) immunostained for the postsynaptic scaffolding protein SHANK3 (cyan) and the vesicular Glutamate transporter vGlut1 (yellow). Left panel shows dendrite of Parvalbumin (PV) positive neuron, right panel shows dendrite of a nonPV neuron. Arrows show SHANK3 and vGlut1 colocalizing puncta. Scale bar, 10 μm. **D**: Quantification of 14 and 12 cells, respectively (N = 2 independent cultures) stained as in (**C**) to determine the degree of colocalisation between SHANK3 and vGlut1. **E**: Fixed hippocampal neurons immunostained against PSD95, PV neuron (upper panel) and nonPV neuron (lower panel) depicted here. **F**: Quantification of PSD95 puncta as shown in (**E**). **G**: tauSTED images of dendrite of PV cell immunostained for α-tubulin and SHANK3. Scale bar, 10 µm. Zoom ins indicated shown in the panel right. **H**: Dissociated hippocampal neuron culture (DIV15) immunostained for the postsynaptic scaffolding protein SHANK3 (magenta) and microtubule minus end binding protein CAMSAP2 (green). Upper panel shows dendrite of Parvalbumin (PV) positive neuron, lower panel shows dendrite of a nonPV neuron. Arrows show SHANK3 and CAMSAP2 colocalising puncta. Scale bar, 10 μm. **I**: Quantification of 16 and 17 cells, respectively (N = 2 independent cultures) stained as in (**A**) to determine the degree of colocalisation between SHANK3 and CAMSAP2.

When localised to excitatory shaft synapses, SHANK3 would be in direct proximity not only of the actin cytoskeleton but likely also the MT network of the dendritic lumen including the MT lattice and minus ends that affect stability and dynamics of the whole MT. We first analysed the abundance of SHANK3 at excitatory synapses of PV neurons. DIV15 neurons were immunostained against SHANK3 and the presynaptic, excitatory marker vesicular glutamate transporter 1 (vGlut1) (***Fig***1C-D). We did not find differences in the degree of colocalisation of the pre- and postsynaptic excitatory proteins in PV neurons as compared to excitatory neurons, suggesting that SHANK3 is also a major PSD scaffold in both neuronal types.

When examining the nanostructural organisation of the F-actin cytoskeleton in PV neurons, we noticed an overall larger dendritic diameter (***Fig* EV**1C). Dendrites of PV neurons contained a more prominent layer of cortical actin (***Fig* EV**1D) as well as a lower density of actin patches per 10 µm (***Fig* EV**1E-F). This indicated that the different actin organisation in PV neurons could impact the degree of the synaptic association to the MT network, potentially influencing intracellular transport and impact synapse and dendritic stability. Similar to previous reports, we additionally found that dendrites of aspiny PV neurons contained a higher density of postsynaptic puncta (15 / 10 µm) than spiny hippocampal neurons (8 / 10 µm, ***Fig***1E-F).

To test whether MTs could indeed associate with excitatory postsynapses of PV neurons, we performed super-resolution microscopy. Using super-resolution tauSTED imaging, we took volumetric stacks of dendrites stained against α-tubulin and SHANK3. We found that SHANK3 positive puncta were frequently found in contact with the MT cytoskeleton of PV dendrites (***Fig***1G). This demonstrated that both MT lattice and ends could indeed be in close physical proximity to SHANK3-positive postsynapses. This proximity suggested the possibility of microtubule nucleation or stabilisation occurring at synapses. To find out how MTs associate with SHANK3-containing postsynaptic sites in aspiny PV neurons, we set out to investigate the colocalisation between SHANK3 and CAMSAPs. CAMSAP2 is the most abundant of the microtubule minus-end binding and stabilising protein family CAMSAP in the hippocampus (*25*). In dissociated hippocampal rat neurons immunostained against SHANK3, CAMSAP2 and PV, colocalisation analysis revealed a higher degree of colocalisation in PV neurons than in excitatory neurons (48% vs 30%) (***Fig***1H-I). This was not due to differences in total protein levels as absolute CAMSAP2 and SHANK3 levels did not show significant differences between PV neurons and other neurons (***Fig* EV**1G-H). This indicated that the MT minus end of non-centrosomal MTs could be in contact with scaffolding proteins at the postsynaptic density of PV neurons.

### Dynamic MT plus ends frequently emerge near excitatory synapses of PV neurons

We next investigated how MTs in PV neurons are organised. We imaged dynamic MT plus ends in PV neurons using MacF18 coupled to mRuby2 (***Fig***2A) which binds to the plus ends of growing MTs (*12*). First, we analysed the directionality of MT outgrowth in PV neurons (***Fig***2B-D). We found that approximately 85% of MTs in proximal and medial dendrites of PV neurons grew with their plus end towards the periphery (***Fig***2D, (*7*, *12*)). As a comparison, dendritic microtubules of mammalian, excitatory neurons have been found to exhibit mixed polarity by both live imaging and EM analysis (*7*, *12*, *13*, *50*), the percentage depending on proximity to the soma and the developmental stage (*7*, *12*)

**Figure 2:**
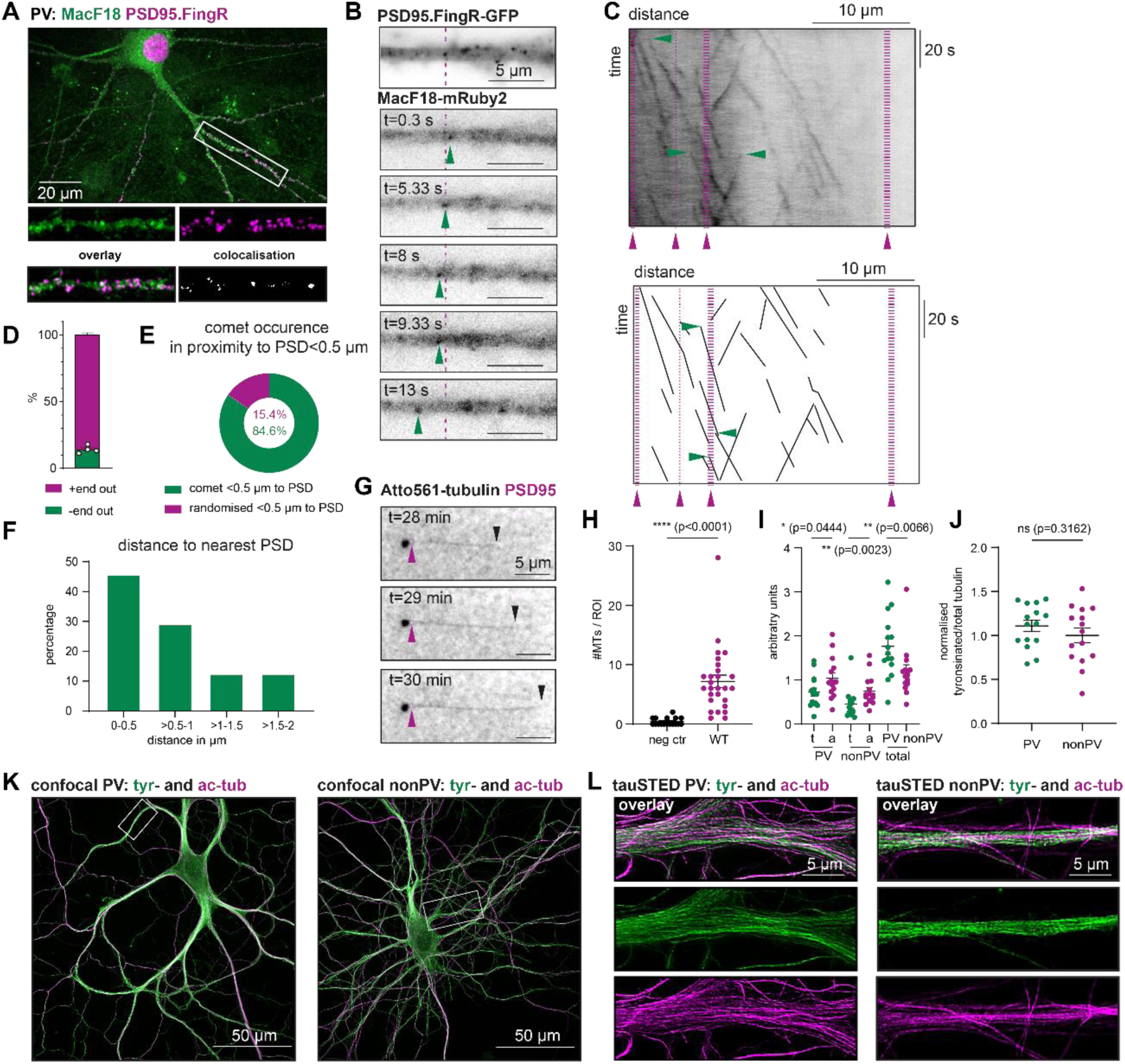
Dynamic MT plus ends emerge close to PSDs of parvalbumin neurons. **A**: Fixed hippocampal neurons labelled with PSD95.FingR-GFP and MacF18-mRuby2, lower panel: overlap of MacF18 and PSD95 signal. **B**: MacF18 emerges in vicinity to PSD on a PV neuron, arrow depicting movement of a selected MacF18 comet. **C**: Example kymograph of associated hippocampal neuron culture (DIV13 - 22) transfected with the dynamic MT marker MacF18-mRuby2 and PSD95.FingR. Reconstruction of kymograph as shown in the lower panel. Arrows in purple show positions of PSDs, arrows in green depict emergence of MacF18 comets. **D**: Polarity quantification of 5 independent cultures (22 dendrites of 12 cells). **E**: Analysis of MacF18 comets emerging in the proximity to PSD95.FingR+ spots normalised to a random distribution of emerging MacF18 comets. **F**: Histogram depicting distance to PSD at which MacF18 comets emerge. **G**: Representative microtubule growth in reaction chamber containing immobilised wildtype synaptosomes from adult mice. Black arrow shows growing plus end of a MT, arrow in magenta indicates PSD95. **H**: Quantification of data as shown in (**G**). Number of microtubules per ROI. **I**: Quantification of data as shown in ***S***2D-E. t = tyrosinated tubulin, a = acetylated tubulin, total = tyrosinated + acetylated tubulin. **J**: Ratio of data as shown in (**I**), normalised to nonPV ratio of tyrosinated/total tubulin. **K-L**: Deconvolved confocal z-stacks of dissociated hippocampal PV and nonPV neurons (**K**) and single planes of tauSTED images of PV (**L**, left panel) and nonPV (**L**, right panel) stained for tyrosinated and acetylated tubulin.

Next, we quantified the distance between the emergence of MacF18 comets and nearby postsynaptic densities (PSDs) labelled with a GFP-tagged PSD95.FingR that binds endogenous PSD95. Due to the particularly high postsynaptic puncta density on PV neurons (***Fig***1E-F), as a proof-of-principle we focused on neurites with a comparatively lower density of PSDs to allow for the observation of MT growth emerging in proximity to PSDs (***Fig***2B). We frequently found dynamic MTs emerging near PSDs (***Fig***2B) in the range of up to 0.5 µm distance when compared to a random distribution of emerging comets (***Fig***2E). In a range of 0-2 µm, we saw that most comets emerged within 0-0.5 µm distance to the closest PSD (***Fig***2F). We further found that PSD95 puncta had higher gamma-tubulin levels in PV neurons (***Fig* EV**2A-B). This suggests that synapses of PV neurons are in proximity to gamma-tubulin, which is one of the major MT nucleation factors (*51*, *52*).

To more directly confirm that this emergence of comets was related to the presence of excitatory synapses in their vicinity, we set up an *in vitro* system in which we immobilised fractionated synaptosomes. To that end, we used synaptosomes purified over a sucrose gradient and differential centrifugation steps. If synaptosomes were to provide MT nucleation promoting factors, it would allow us to observe MT growth in samples supplied with synaptosomes and not in control samples in which a low tubulin concentration of 15 µM was used that was insufficient to cause MT nucleation (***Fig***2G-H). We saw that through the addition of synaptosomes, MT growth was indeed induced (***Fig***2G-H).

In an earlier study by (*6*), it was shown that MT bundles with more stable MTs (acetylated and detyrosinated) consist of minus-end-out orientation, whereas more dynamic ones (tyrosinated) have their plus-ends directed away from the soma. How the overall prevalence of plus-end-out MTs and the initiation of MT growth at the PSD influence the distribution of stable and dynamic MTs in dendrites of PV neurons remains an open question. We thus compared the distribution of tyrosinated to acetylated MTs to the known distribution within dendrites of excitatory neurons both with confocal and tauSTED imaging. We first compared levels of acetylated and tyrosinated MTs within and between PV and nonPV neurons. We found that similarly to excitatory neurons, dendrites of PV neurons contained tyrosinated and acetylated MTs (***Fig***2I, K-L, ***Fig* EV**2D-G). However, we could also see that the total amount of MTs per area was significantly higher in PV neurons compared to nonPV neurons. However, when comparing the ratio of dynamic and stable MTs we saw no clear difference between PV and nonPV neurons (***Fig***2J). Acetylated MTs were arranged in bundles in both cell types (***Fig***2J). Next, we analysed the distribution of stable and dynamic MTs by plotting line profiles and analysing the area under the curve. We did not find differences between PV and nonPV neurons. PV and nonPV neurons showed a ratio of 0.97 and 0.95 acetylated/tyrosinated tubulin, respectively (***Fig* EV**2F-G). This in turn indicated that the presence of MT plus ends pointing to the periphery of dendrites in PV neurons could not be explained by a differering ratio of stable to dynamic MTs. We thus hypothesised that MT growth events emerging in close proximity to a PSD could be emerging from MT seeds stabilised by CAMSAP2, which serve as MT outgrowth sites.

### CAMSAP2 interacts with the synaptic scaffolding protein SHANK3

To gain mechanistic insights into how CAMSAP2 associates with the PSD, we performed a mass spectrometry screening of GFP-CAMSAP2-associated protein complexes enriched from rat brain extracts using affinity purifications. We found known CAMSAP2 interactors such as WDR47 (WD Repeat Domain 47) and spectrin. A comparison with the GeneOntology database “Biological Process 2023 Analysis” revealed CAMSAP2 interactors were commonly annotated as playing roles in cytoskeletal protein binding or were structural constituents of synapses (***Fig***3A). Indeed 74 of 212 identified CAMSAP2 interactors could be identified with the synaptic database SynGO (***Fig***3B) including SHANK3 and spectrin (SPTBN2). This finding led us to hypothesise that there could in fact be an interaction with postsynaptic proteins. Additionally, we could show that most of the interactors found in the CAMSAP2 mass spectrometry screen were also detected in the SHANK3 mass spectrometry screen (***Fig* EV**3A). Of these overlapping interactors, 39 were found to be synaptic (***S***3B). Overlapping hits such as MAP2 and MacF1 were also found to be annotated as involved in cytoskeletal organisation (***S***3C). This further indicated a role for CAMSAP2 in connecting the PSD to the MT cytoskeleton.

This interaction could be confirmed through endogenous co-immunoprecipitations (coIPs) of proteins from brain lysate with CAMSAP2 in which CAMSAP2 from rat brain extract was used as bait and the different SHANK proteins were detected as prey (***Fig***3C). In the IP fraction, CAMSAP2 was detected and it was absent from all negative controls. In the coIP fractions as well as in the input, SHANK2 and 3 could be detected (***Fig***3C). SHANK1 could only be found in the input sample, but not in the coIP fraction and was the only SHANK-family member that did not interact with CAMSAP2 (***Fig***3C).

Next, we asked whether the binding was direct and where the binding regions on CAMSAP2 and SHANK3 were located (***Fig***3D-E). To that end, we pulled down GFP-fused truncations of CAMSAP2 at (***Fig* EV**3D) and incubated them with rat brain lysate. Immunoblotting against SHANK3 indicated that the interaction was mediated through the MT binding domain (MBD) of CAMSAP2 (***Fig***3F, H). Similarly, the binding site of CAMSAP2 on SHANK3 was mapped (***Fig***3G, I). Truncations of SHANK3 (1–676) containing the domains: SPN, ARR, PDZ and SH3 domain were pulled down from HEK293T cells and incubated with the extract of HEK293T cells overexpressing full length (FL) CAMSAP2 (***Fig***3G, I). CAMSAP2 could be detected in all samples containing either the SPN or the ARR domain of SHANK3 but not in those just containing the SH3 domain or the negative control (***Fig***3G, I). This analysis further confirmed the binding and determined the ARR and SPN domain of SHANK3 to interact with CAMSAP2. The MBD domain, which anchors CAMSAP2 to the MT lattice, was also interacting with SHANK3, whereas the second MT-binding domain, the CKK, was not involved and remained accessible for MT association.

To find out whether SHANK3 and MTs would competitively or cooperatively bind to the MBD, we performed an *in vitro* co-pelleting assay using purified proteins. MTs were polymerised and CAMSAP2 MBD and additionally SHANK3 (***S***3E) was added in selected samples. Following ultracentrifugation, supernatant and pellet samples were collected, and protein levels analysed using immunoblotting. We found that CAMSAP2 MBD pelleted in the presence of MTs, but MBD pelleting was decreased when SHANK3 was additionally present (***Fig***3J). CAMSAP2 FL pelleted in the presence of MTs, and was able to pellet in the presence of SHANK3. This indicates that SHANK3 and MTs cannot bind to the MBD simultaneously, but when the MBD is occupied by SHANK3 the CKK domain of CAMSAP2 might still be available for the association with the MT minus ends as evidenced by a less severe reduction of CAMSAP2 FL pelleting in the presence of SHANK3. This result was especially intriguing since CAMSAP2 is known to preferentially bind to the minus end. CAMSAP2 can bind to MTs with two different domains: a C-terminal domain present in all three CAMSAPs responsible for the MT minus binding behaviour (CKK) and another microtubule binding domain (MBD) involved in the association with the MT lattice. We thus propose that CAMSAP2 could bind to SHANK3 while bound to MTs thus anchoring MTs at the PSD (***Fig***3K).

**Figure 3:**
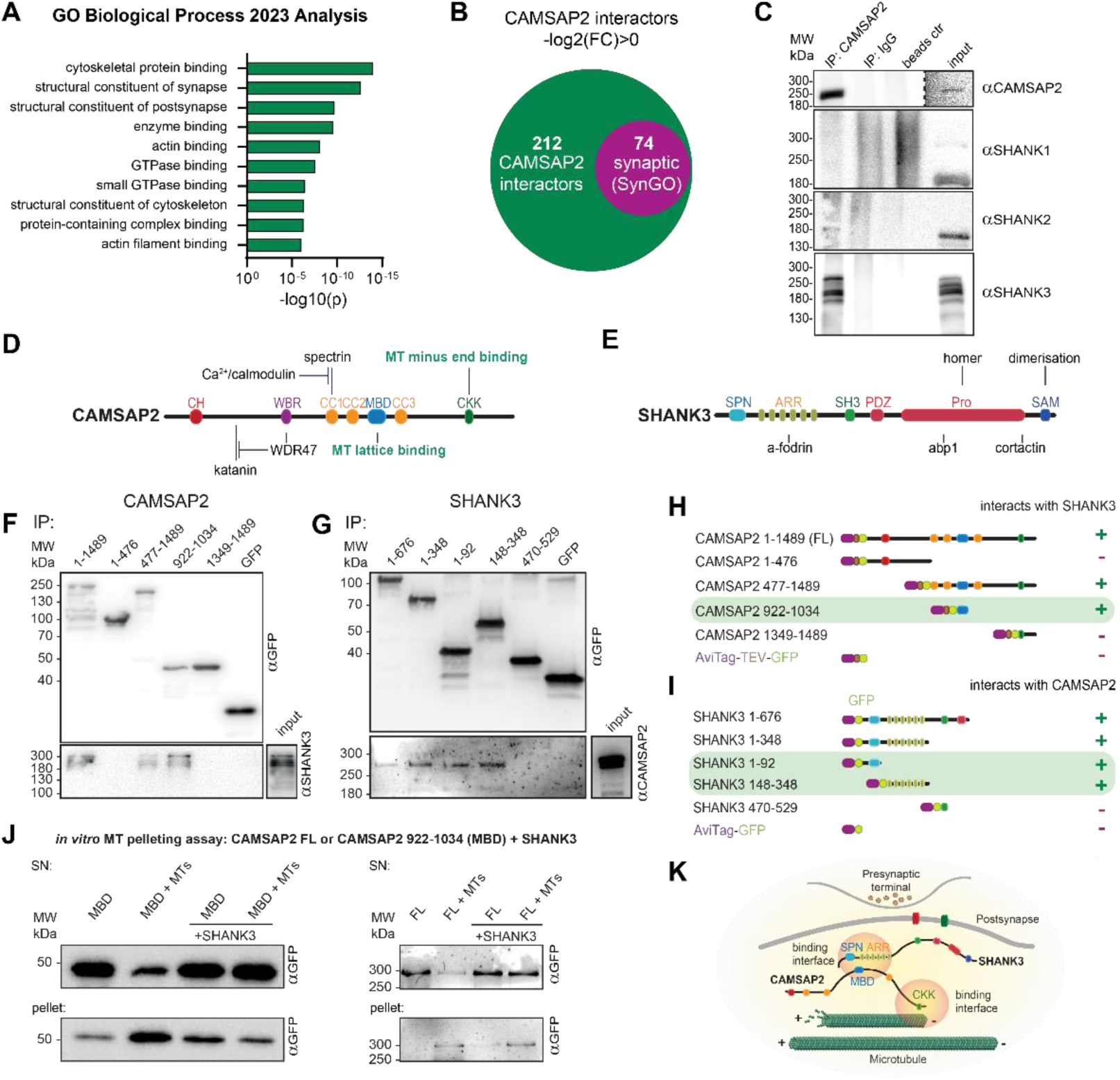
CAMSAP2 interacts with SHANK3 through its microtubule binding domain. **A**: GO Biological Process analysis of mass spectrometry data of a CAMSAP2 pulldown. **B**: Total number and share of synaptic proteins found in the CAMSAP2 pulldown mass spectrometry screen through a comparison with the SynGO database. **C**: coIP of CAMSAP2 and SHANK3 from rat brain lysate using an antibody raised against CAMSAP2 cross-linked to Protein A&G. Detected using antibodies raised against SHANK3 and CAMSAP2. From left to right: CoIP sample, IgG control (brain lysate incubated with donkey anti-goat antibody), control (brain lysate incubated with Protein A+G mixture) and input. **D**: Graphic summary of CAMSAP2 structure and proteins interacting at specific domains. **E**: Graphic summary of SHANK3 structure and proteins interacting at specific domains. **F**: Example immunoblot of pulldown samples of CAMSAP2 truncations overexpressed in HEK293T cells, co-precipitated with CAMSAP2-FL from HEK cell lysate. N = 3 **G**: Example immunoblot of pulldown samples of SHANK3 truncations overexpressed in HEK293T cells, co-precipitated with CAMSAP2-FL from rat brain lysate. N = 4 **H**: CAMSAP2 truncations domain composition and summary of binding to CAMSAP2 from data as shown in (**F**). **I**: SHANK3 truncations domain composition and summary of binding to CAMSAP2 from data as shown in (**G**). **J**: Representative immunoblot showing supernatant (SN) and pellet samples of CAMSAP2 MBD (left panel) or FL (right panel) obtained in a co-pelleting assay. **K**: Schematic showing CAMSAP2-SHANK3 binding interface while attached to the postsynaptic density and CAMSAP2-microtubule binding interface.

### CAMSAP2 MBD reduces dendritic arbour length

Having discovered that the CAMSAP2 MBD could bind to either MTs or to SHANK3, its overexpression would be expected to compete with endogenous CAMSAP2 and be used to decouple SHANK3 from CAMSAP2 and thus detach MTs from the postsynaptic density. We next asked whether this would alter processes affected by MT dynamics such as dendritic branching and dendritic arbour length. Since we found a higher colocalisation of SHANK3 and CAMSAP2 in aspiny PV neurons (***Fig***1H), we would expect an effect to be more pronounced in these neurons.

To test this, we overexpressed the CAMSAP2 MBD in hippocampal rat neurons at DIV7 and explored the effect on dendritic branching at DIV14.

We first investigated this effect in nonPV neurons (***Fig***4A-B). Here, we found that the overexpression of CAMSAP2 MBD decreased the dendritic arbour length (***Fig***4E), but did not significantly alter dendritic branch length (***Fig***4F) or the number of dendritic branches (***Fig***4G). In the corresponding Sholl-analysis (***Fig***4H), we found that the area under the curve was 709 µm^2^ for CAMSAP2 MBD and 895 µm^2^ for GFP overexpression, further highlighting the overall change in dendritic branching.

**Figure 4:**
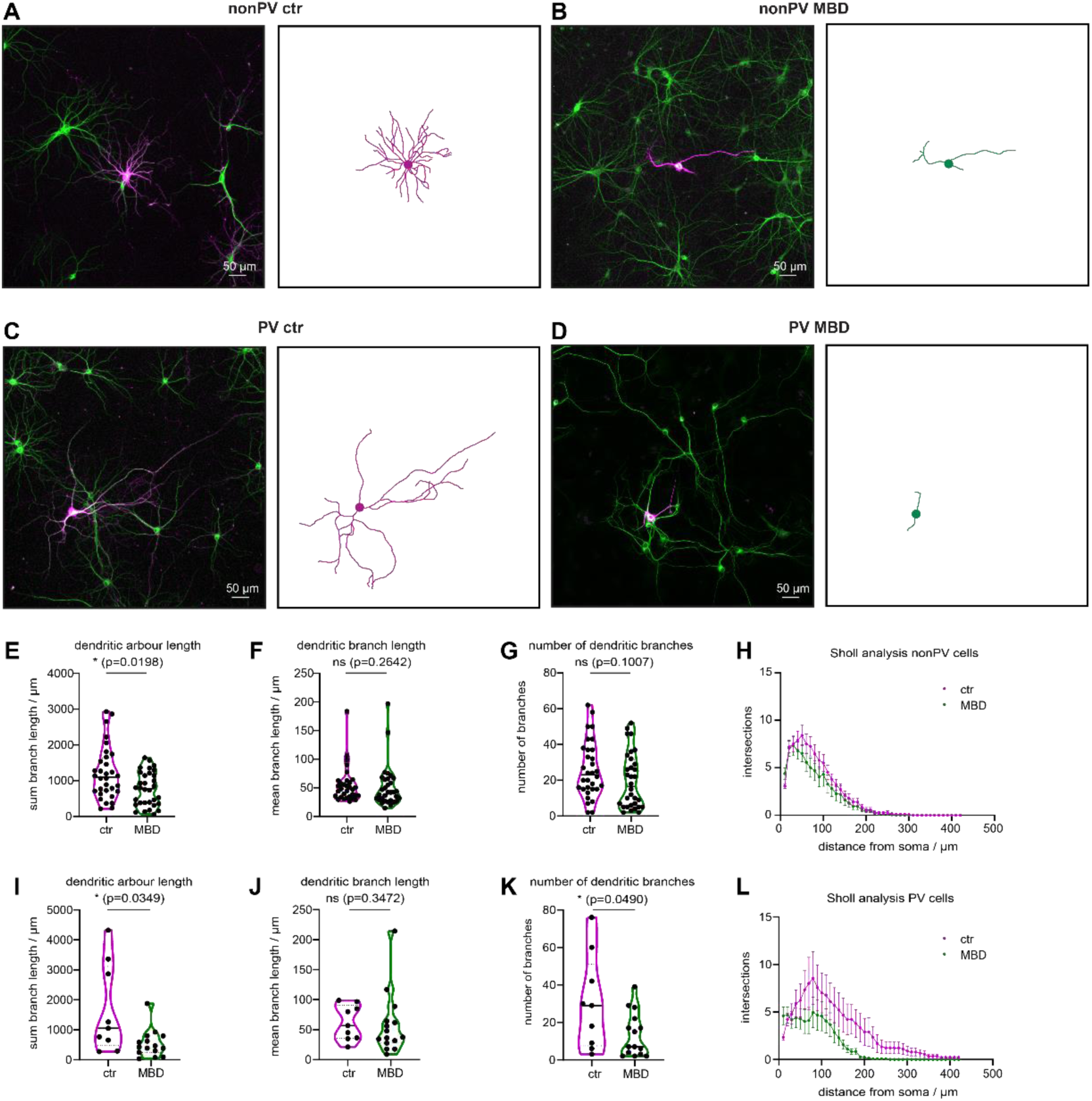
CAMSAP2 MBD overexpression reduces dendritic arbour length. **A-B**: Representative images and traced nonPV neurons expressing GFP (ctr) (**A**) or CAMSAP2 MBD (**B**) using SNT, soma added post-analysis. Scale bar, 50 µm. **C-D**: Representative images and traced PV neurons expressing GFP (ctr) (**C**) or CAMSAP2 MBD (**D**) using SNT, soma added post-analysis. Scale bar, 50 µm. **E-G**: Quantifications of traces as shown in (**A-B**) using SNT. Total dendritic arbour length (**E**), length of individual branches (**F**) and number of dendritic branches (**G**) compared between WT and SHANK3_L68P_ neurons. **H:** Sholl analysis of traces as shown in (**A-B**) using SNT. **I-K**: Quantifications of traces as shown in (**C-D**) using SNT. Total dendritic arbour length (**C**), length of individual branches (**D**) and number of dendritic branches (**E**) compared between WT and SHANK3_L68P_ neurons. **L:** Sholl analysis of traces as shown in (**C-D**) using SNT.

To see whether the increased colocalisation of SHANK3 and CAMSAP2 in PV neurons would lead to an increased susceptibility to disrupting their interaction, we then investigated the effect of the expression of CAMSAP2 MBD in PV neurons (***Fig***4C-D). Here, we found a stronger, more profound effect of the expression of CAMSAP2 MBD. The overexpression of CAMSAP2 MBD decreased dendritic arbour length (***Fig***4I), did not significantly reduce dendritic branch length (***Fig***4J), but reduced the number of dendritic branches (***Fig***4K), highlighting the susceptibility of PV neurons to the disruption of CAMSAP2-SHANK3 binding, which was an effect only observed as a trend in nonPV neurons. In the corresponding Sholl-analysis (***Fig***4L), we found that the area under the curve was 601 µm^2^ for CAMSAP2 MBD and 1275 µm^2^ for GFP overexpression, clearly displaying the altered dendritic phenotype of PV neurons. Especially at a distance of >150 µm from the soma, the different number of intersections became very clear to observe, with CAMSAP2 MBD-expressing neurons having fewer intersections (***Fig***4L).

This indicated a clear dendritic phenotype caused by the expression of the CAMSAP2 MBD with a more pronounced effect in PV neurons, likely caused by the higher base level-association of CAMSAP2 and SHANK3.

### ASD-associated SHANK3_L68P_ mutation enhances interaction with CAMSAP2

We next focused on the SPN and ARR domains of SHANK3 that bind to CAMSAP2 (***Fig***3G, I). Previously, we have shown that the intramolecular interaction between these two domains is differentially altered in the ASD-associated SHANK3_R12C_ and SHANK3_L68P_ mutants (*39*). We thus set out to see whether these mutations would influence the binding to CAMSAP2.

First, to get a broader view on the impact of both mutations on the SHANK3 interactome, we performed a mass spectrometry screening on samples obtained in the same way as SHANK3 truncations which were overexpressed for the determination of the CAMSAP2-binding region, this time including the ASD-associated SHANK3_R12C_ and SHANK3_L68P_ mutations. We found that the mutations resulted in loss- or gain of function interactions with a multitude of MT cytoskeleton associated proteins including CAMSAPs, MAPs and MT based motors such as KIF7 (***Fig***5A, B). Interestingly, CAMSAP2 showed increased binding to SHANK3_L68P_ but not to SHANK3_R12C_. The SHANK3 interactor SHANK1 was found to show increased binding to both mutants. Therefore, in a validation experiment we next overexpressed amino acid 1-676 truncations of SHANK3 WT or L68P mutation in HEK293T cells and then performed a coIP using the GFP-tagged SHANK3 truncations and adult rat brain lysate containing CAMSAP2 was added. In both SHANK3 WT and L68P samples, CAMSAP2 was detected with protein levels showing an average five-fold increase in the L68P mutant samples, (***Fig***5C-D). This indicates that the changes in intramolecular folding of SHANK3 caused by single point mutations had vast effects on its interactome and could influence the MT cytoskeletal properties of neurons.

**Figure 5:**
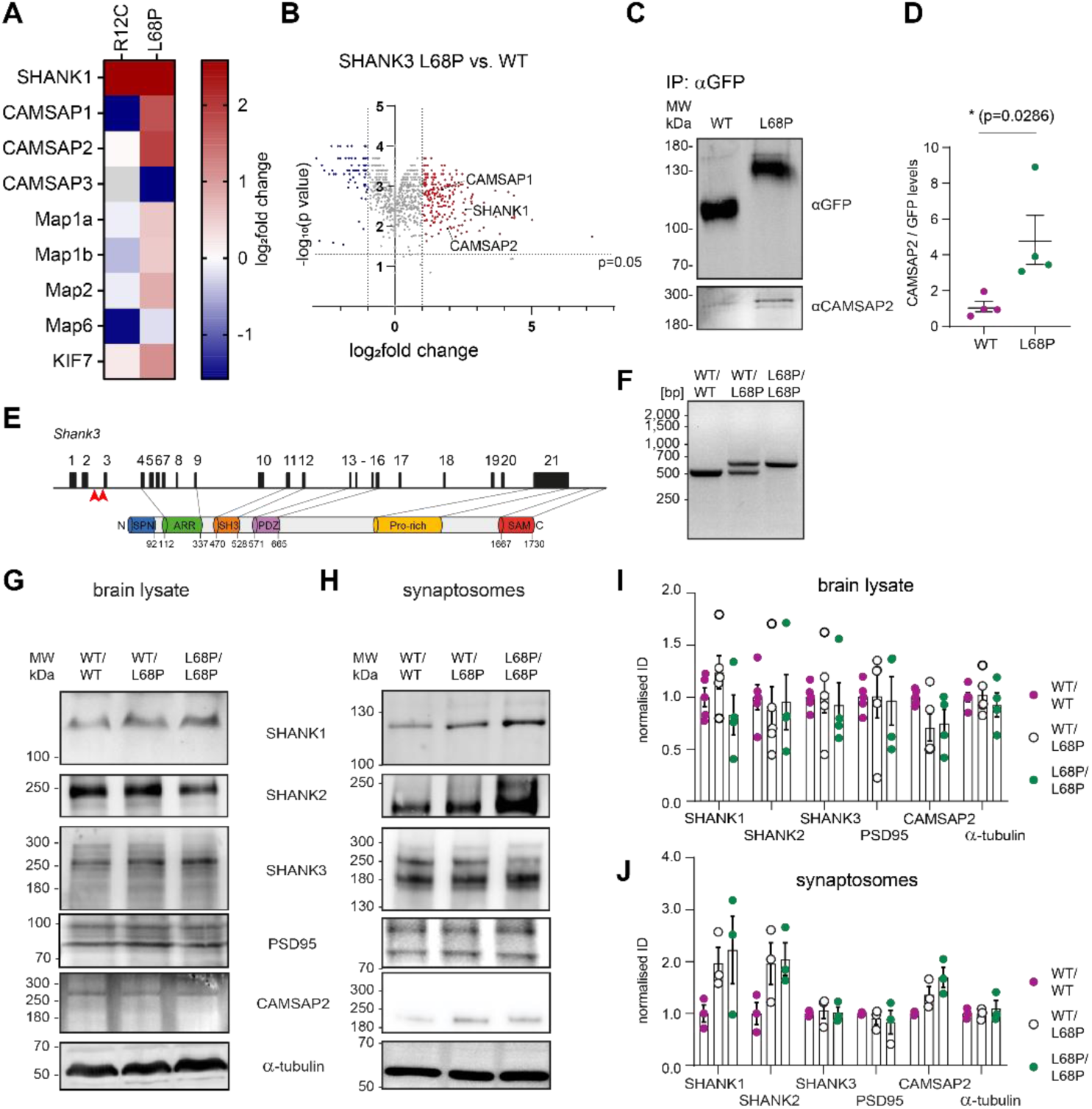
ASD-associated SHANK3 L68P mutation increases interaction with CAMSAP2. **A**: Heatmap showing log_2_fold change of SHANK3 point mutations on selected SHANK3 interaction partners in pull-down experiments. **B**: Volcano plot showing the impact of the SHANK3_L68P_ mutation on the CAMSAP2 protein levels. PEP value/10000 used for log_10_ of p value. **C**: Representative immunoblot of pulldown of SHANK3 1-676 WT or L68P and subsequent protein binding from rat brain lysate. using antibodies raised against SHANK3 and CAMSAP2. N = 4 animals. From left to right: coIP sample WT and L68P. **D**: CAMSAP2 levels normalised to SHANK3 input. **E**: Schematic depicting the organisation of the gene encoding SHANK3. **F**: Genotyping of mouse line encoding SHANK3_L68P_. **G-H**: Representative immunoblots of mouse brain lysates (**G**) or synaptosomes (**H**) of SHANK3_L68P_ KI mice with antibodies against CAMSAP2, SHANK3 and PSD95. From left to right: wildtype brain lysate, heterozygous, homozygous KI samples. 30 μg protein per lane. **I-J**: Quantification of brain lysate samples (**C**) or synaptosome samples (**D**) using the gel analysis function of FIJI. Normalised CAMSAP2 levels and ratios of CAMSAP2 levels and SHANK3 levels depicted as mean +-SEM. Normalised to ratios of corresponding WT values from the same preparation.

Next, we moved to a genetically modified mouse line carrying SHANK3 with an L68P mutation knock-in (KI, ***Fig***5E-F). First, we characterised protein levels of the SHANK3_L68P_ KI mouse line by comparing protein levels from wild type and mutants in brain lysates and subcellularly fractionated synapses (***Fig***5G-J) since we were interested in an interaction that is expected to predominantly take place at the postsynaptic density. Protein levels of selected proteins that showed a gain-of-function in SHANK3_L68P_ mutation such as SHANK1, SHANK3, PSD95, CAMSAP2 and α-tubulin were similar across genotypes in brain lysate samples (***Fig***5G-I). SHANK2 protein levels were slightly reduced, compared to wildtype over hetero- to homozygous SHANK3_L68P_ KI samples. In SHANK3_L68P_ KI synaptosomes SHANK1, SHANK2 and CAMSAP2 levels showed trends of being increased in hetero- and homozygous SHANK3_L68P_ KI samples in gene-dose dependent manner while SHANK3, α-tubulin and PSD95 levels were constant across genotypes (***Fig***5H-J). This indicates that the L68P mutation altered synaptic protein levels of SHANK3 interactors including CAMSAP2, likely due to gain-of-function interactions.

### Synaptic SHANK3 associates with MT seeds and the L68P mutation enhances this association

To investigate the relation between the postsynaptic density containing SHANK3_WT_ or SHANK3_L68P_ and MTs, we used a modified version of our established *in vitro* approach to immobilise fractionated synaptosomes (***Fig* EV**6A) that were then allowed to associate with HyLite tubulin (***Fig***6A). Synaptosomes were stained with pre-labelled PSD95 nanobodies (***Fig***6A). This labelling was possible due to the postsynaptic density often being accessible following synaptosome-preparation. We then added HyLite tubulin and observed tubulin association to synaptosomes after ∼30-60 min. This allowed us to observe tubulin structures including short “MT seeds” (***Fig***6B). We then compared whether the SHANK3_L68P_ mutation altered the association between the PSD and tubulin. We found that around 10% of MTs colocalised with labelled synaptosomes in the wildtype and SHANK3_L68P_ KI synaptosomes (***Fig* EV**6B-C). When quantifying the degree of colocalisation of WT synaptosomes with tubulin, we found around 7% of synaptosomes had MT seeds associated with them (***Fig***6C). In synaptosomes obtained from homozygous SHANK3_L68P_ KI mice, around 35% of synaptosomes had MT seeds associated with them (***Fig***6C). This indicated that MT seeds were increasingly stabilised and anchored at postsynaptic densities of synapses carrying the SHANK3_L68P_ mutation. We again used the *in vitro* assay optimised for visualising MT growth (***S***2C). Here, we found that more MTs grew in growth chambers containing SHANK3_L68P_ synaptosomes (***Fig***6D). Note that the number of MTs in both datasets are normalised to the average of labelled synaptosomes found in the growth chambers. Wildtype data already shown from ***Fig***2H was used to calculate this normalisation (***Fig* EV6**D). In our *in vitro* experiment, we focused on comparing homozygous SHANK3_WT_ synapses and homozygous SHANK3_L68P_ synapses since in heterozygous animals, we would not have a way to identify whether fewer synapses contain SHANK3_WT_ or whether the levels of SHANK3_WT_ are reduced in all synapses.

**Figure 6:**
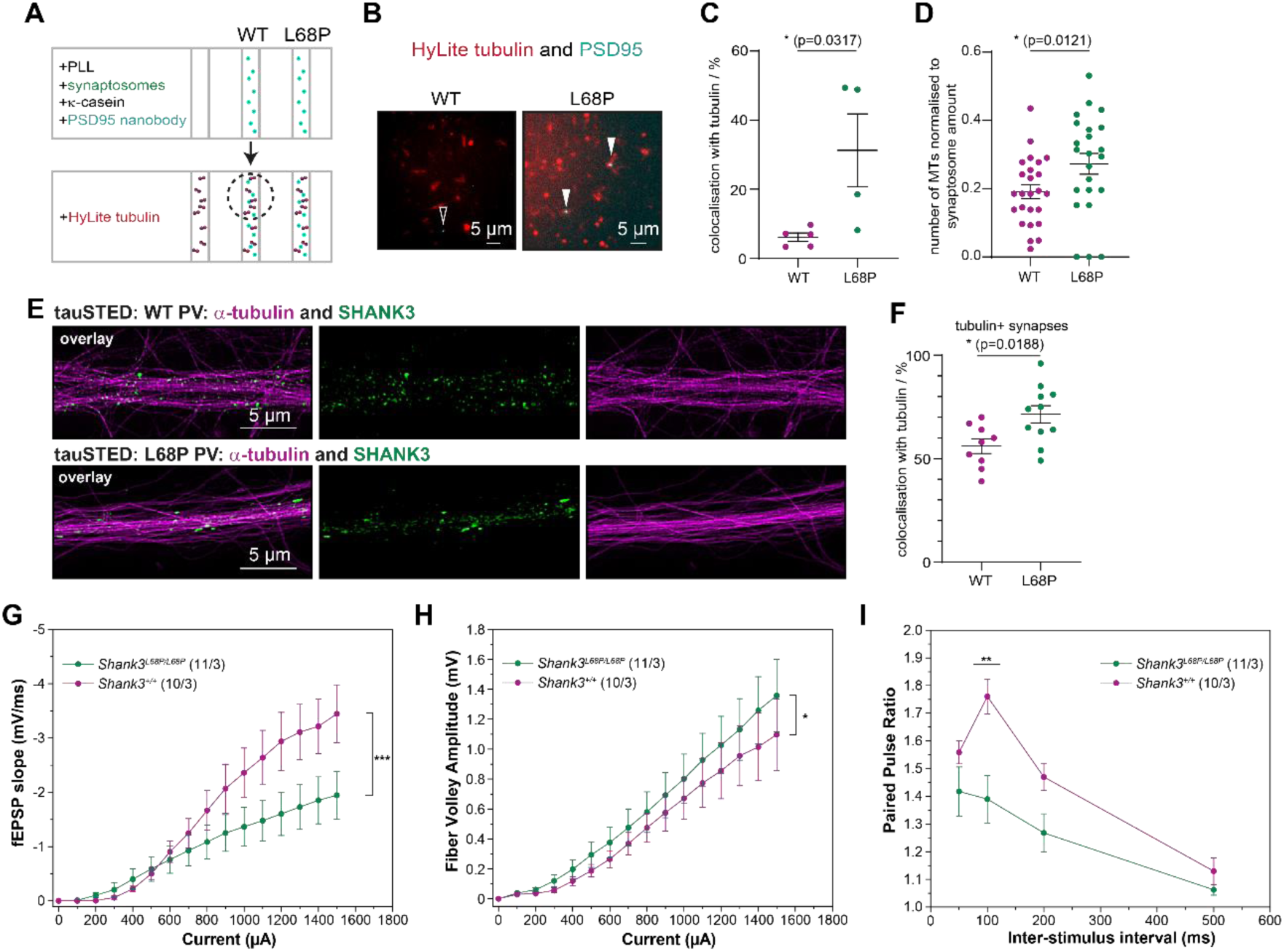
SHANK3_L68P_ mutation increases association with microtubules in parvalbumin neurons. **A**: Schematic showing the workflow of immobilising and labelling synaptosomes on glass slides **B**: MT seeds in relation to WT and SHANK3_L68P_ synaptosomes. Colocalisation marked with filled arrows, synaptosome not colocalising with tubulin marked with empty arrow. **C**: Number of synaptosomes with associated MT seeds as in (**B**). **D**: Quantification of data as shown in Figure 2G-H including SHANK3_L68P_ synaptosomes. Number of microtubules per ROI, normalised to the average number of synaptosomes per reaction chamber. **E**: WT (upper panel) and SHANK3_L68P_ (lower panel): Dissociated hippocampal neuron culture (DIV15) immunostained for the postsynaptic scaffolding protein SHANK3 (green) and α-tubulin (magenta). Scale bar, 5 μm. **F**: Quantification of 9 and 11 cells, respectively (N = 2 independent cultures) stained as in (**E**) to determine the degree of colocalisation between SHANK3 and α-tubulin. **G**: Input-output curves measured on acute slices (male, 2 - 4 months old) (two-way repeated measures ANOVA with Tukey posthoc test, *** p < 0.0005). Number of slices and animal (N) indicated in the inlet. **H**: Fibre volley amplitude (two-way repeated measures ANOVA with Tukey posthoc test, * p < 0.05). **I**: Paired pulse ratio (two-way repeated measures ANOVA with Tukey posthoc test, overall: p(*Shank3_WT_/Shank3_L68P_*) < 0.0005). Inter-stimulus interval of 100 ms (two-way repeated measures ANOVA with Tukey posthoc test, genotype within inter-stimulus interval = 100 ms: p(*Shank3_WT_/Shank3_L68P_*) < 0.0005).

To follow up on these *in vitro* results, we then investigated the association of dendritic MTs with SHANK3 in PV neurons containing SHANK3_WT_ or SHANK3_L68P_. Consistently, in tauSTED images of DIV15 neurons of the L68P KI mouse line (homozygous), we found that SHANK3 puncta showed a higher degree of colocalisation with tubulin in the homozygous L68P condition (∼72%) than in the homozygous WT condition (∼56%) (***Fig***6E-F). This was not caused by SHANK3 puncta occupying a larger area in the SHANK3_L68P_ condition as cluster sizes were comparable between the two conditions (***Fig* EV**6E).

Synaptic transmission and plasticity require constant remodelling and molecular transport of cargo to and from the synapse. Alterations in polarity and MT network accessibility may therefore broadly modulate network activity by shaping PV neuron excitability and firing rates. We therefore investigated whether the SHANK3_L68P_ mutation impacts hippocampal network activity. *Ex vivo* electrophysiological recordings were performed using acute hippocampal slices from 2-4 month old SHANK3_L68P_ KI mice and SHANK3_WT_ littermates. Action potentials were evoked in the Shaffer collateral (SC) fibres originating from the CA3 region while field excitatory postsynaptic potentials (fEPSP) were recorded in the dendritic area of CA1. We assessed excitatory synaptic transmission at SC-CA1 pyramidal cells synapses by measuring the amplitude of the fibre volley (presynaptic input) and the fEPSP slope (postsynaptic response). Hippocampal slices from SHANK3_L68P_ KI mice exhibited an increased fibre volley amplitude, indicating enhanced presynaptic fibre recruitment, plausibly caused by a decrease in effective PV neuron inhibition on excitatory neurons. However, a markedly diminished fEPSP slope was observed in SHANK3_L68P_ KI mice, indicating a decreased postsynaptic response (***Fig***6G-H). Next, we assessed short-term plasticity at SC–CA1 synapses by measuring the paired-pulse ratio (PPR) following two stimuli delivered at 50, 100, 200 and 500 ms interstimulus intervals. This form of short-term synaptic plasticity is determined, at least in part, by changes in probability of neurotransmitter release. We found that the PPR was reduced in SHANK3_L68P_ hippocampal slices compared to WT slices (***Fig***6*I*), suggesting an increased presynaptic vesicle release probability at glutamatergic synapses and/or a reduction in functional postsynaptic sites.

Since both length and branching of the dendritic arbour are determined by MT stability, we next tested whether PV neurons containing SHANK3_L68P_ showed differences in their dendritic organisation and how the depletion of CAMSAP2 affected it additionally. To that end, SHANK3_L68P_ or SHANK3_WT_ neurons were transduced at DIV7 to express Cre under a promotor specifically active in PV neurons. Floxed CAMSAP2 shRNA or scrambled shRNA was then introduced by an AAV, making only cells transduced with both viruses green fluorescent indicating CAMSAP2 KD or a control in PV neurons. We did not use MAP2 to trace dendrites since MAP2 levels were reduced in CAMSAP2 KD PV neurons (***Fig* EV**7), consistent with literature suggesting early CAMSAP2 depletion could reduce MAP2 levels (*25*).

**Figure 7:**
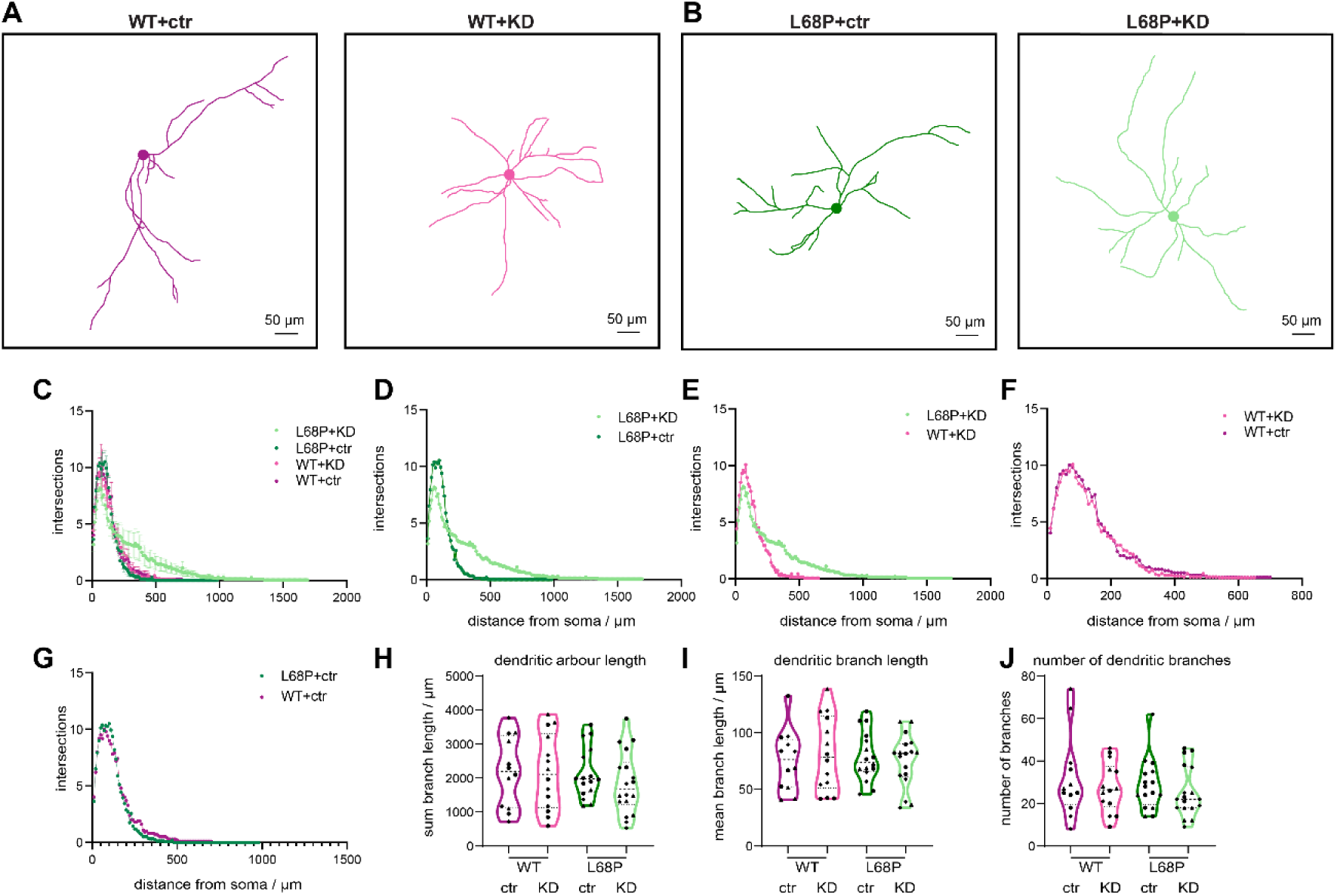
CAMSAP2 depleted SHANK3_L68P_ PV neurons show changed complexity of the dendritic arbour. **A-B:** Representative images and traced SHANK3**_WT_** PV neurons (**A**) or SHANK3**_L68P_** PV neurons (**B**) transcribing scrambled shRNA or CAMSAP2 KD shRNA. Soma added post-analysis. Scale bar, 50 µm. **C:** Sholl analysis of traces as shown in (**A-B**) using SNT. **D-G**: Curves of Sholl analyses as seen in (**C**) displayed without error bars to enhance visibility. **H-J**: Quantifications of traces as shown in (**A-B**) using SNT. Total dendritic arbour length (**H**), length of individual branches (**I**) and number of dendritic branches (**J**) compared between WT and SHANK3_L68P_ neurons transcribing scrambled shRNA or CAMSAP2 KD shRNA.

We found that the Sholl-curve and the area under the curve of SHANK3_L68P_ neurons treated with a CAMSAP2 KD differed from all other groups (***Fig***7C). For better comparability of individual groups, we displayed Sholl-curves in individual graphs without ±SEM (***Fig***7D-G). SHANK3_L68P_ PV neurons in which CAMSAP2 was knocked down showed an increased area under the curve when compared to SHANK3_L68P_ PV neurons with a scrambled control (2391 to 1646 µm^2^) (***Fig***7C-D). Knocking down CAMSAP2 in SHANK3_WT_ PV neurons did not show the same alteration of the dendritic morphology (1659 to 1761 µm^2^) (***Fig***7C, E). The dendritic morphology of SHANK3_WT_ PV neurons in which CAMSAP2 was knocked down was similar to those in which a scrambled control was expressed (***Fig***7C, F). SHANK3_WT_ and SHANK3_L68P_ PV neurons were similar in their dendritic morphology (**Fig**7C, G).

When comparing dendritic arbour length (***Fig***7H), dendritic branch length (***Fig***7I) and number of dendritic branches (**Fig**7J), we found no significant differences between any of the groups.

Altogether, this indicates that CAMSAP2 depletion strongly affects MT cytoskeleton dynamics in SHANK3_L68P_ PV neurons which may be more susceptible to alterations due to the increased interaction between SHANK3_L68P_ and CAMSAP2.

## Discussion

In this work, we found that the MT minus end binding protein CAMSAP2 interacts with the postsynaptic scaffolding protein SHANK3, enabling the association of MT minus ends with the excitatory postsynapse. We find that this interaction is especially of high relevance in PV neurons since they receive their excitatory inputs through synapses located directly on the dendritic shaft. Indeed, an overexpression of the dominant negative CAMSAP2 truncation could decouple CAMSAP2 and MTs from SHANK3, resulting in a dramatic simplification of dendritic architecture. Along the same lines, an ASD-associated mutation of the excitatory postsynaptic density protein SHANK3, resulting in an amino acid exchange from lysin to proline at the position 68, which enhances the binding between SHANK3 and CAMSAP2, also impaired the dendritic architecture of PV neurons obtained from SHANK3_L68P_ mice when simultaneously reducing CAMSAP2 levels. This combined effect highlights the susceptibility of the dendritic MT cytoskeleton of PV neurons in the mutant condition.

Scaffolding proteins are known to connect to the actin cytoskeleton (*53*). In the case of SHANK3, these interactions also entail multiple actin binding proteins such as α-fodrin and cortactin (*54*). In excitatory neurons, a small proportion of glutamatergic synapses is located directly on the dendritic shaft and their PSD is surrounded by a dense mesh of actin filaments devoted of MTs (*55*). We found that PV neurons have fewer actin patches and at the same time, a higher degree of colocalisation with MTs at excitatory synapses (***Fig***1). This could cause changes in cargo supply to synapses e.g. in response to plasticity-inducing stimuli. Furthermore, we found that in contrast to nonPV neurons, the majority of dendritic MTs in PV neurons have their plus end directed to the periphery. We also see a difference in tubulin levels between PV and nonPV neurons which adds further evidence for their distinctly organised dendritic MT cytoskeleton.

We found that in addition to the MT availability at synapses in PV neurons, more SHANK3 puncta were positive for CAMSAP2 compared to nonPV neurons, placing the MT minus end in close proximity of the PSD. Since MT dynamics can be mediated from the minus end, this synapse-MT colocalisation might influence the stability of not just synapses but the MT network. In accordance, patronin (*Drosophila* CAMSAP1 homologue) has been identified as an important player in maintaining dendritic polarity as well as playing a role in influencing dendritic branching (*56*). While our work focuses on the role of the CAMSAP2 and SHANK3 interaction in anchoring MTs to shaft synapses of PV neurons, we cannot exclude that it could also be relevant for shaft synapses of other neuron types or across different species.

Interestingly, we found that disrupting the interaction between SHANK3 and CAMSAP2 by expressing the CAMSAP2 MBD, led to a more pronounced dendritic alteration in PV neurons, hinting to their susceptibility to changes in the synaptic anchoring of MTs. There was similar trend in nonPV neurons, however, despite much larger n numbers, the differences did not reach significance. Overexpressed CAMSAP2 MBD is present at higher levels than endogenous CAMSAP2 and therefore competes with it to e.g. displace it from the synapse where it is bound to SHANK3. Other than a CAMSAP2 KD, it does not affect overall MT stabilisation in the same way, as the presence of CAMSAP2 MBD is not sufficient for effective MT stabilisation (*25*). We further saw that a CAMSAP2 KD strongly affected the morphology of the dendritic arbour of SHANK3_L68P_ PV neurons even when it was expressed at a later stage of neuronal development that coincides with synaptogenesis, although it did not affect the dendritic arbour of SHANK3_WT_ PV neurons. This indicates that the likely stronger interaction of CAMSAP2 with SHANK3_L68P_ increases the vulnerability of the dendritic arbour of PV neurons to perturbations such as the depletion of CAMSAP2, resulting in a dendritic phenotype.

Using synaptosomes and pulldown experiments, we found that the CAMSAP2 binding regions of SHANK3 were the ones most affected by the ASD-associated L68P mutation that we previously structurally characterised using small-angle X-ray scattering (SAXS) (*39*). The SHANK3_L68P_ mutation consistently increased the association with CAMSAP2. We propose that this is the case since the ARR and SPN domains are more accessible in the SHANK3_L68P_ mutant (*57*). Contrary to our expectation, the increased presence of CAMSAP2 at isolated synaptic compartments containing SHANK3_L68P_ did not coincide with an increased colocalisation with MTs but with that of short MT seeds and an increase in MT growth. CAMSAP2 recognises growing MT minus ends and its binding stabilises them, increasing their resistance to depolymerisation (*58*). The increased interaction with tubulin and the presence of CAMSAP2 at SHANK3_L68P_ synapses might therefore stabilise MT seeds that form the foundation for MT nucleation. However, *in vitro* assays in which we used synaptosomes and pure tubulin, are far less complex and only reproduce a specific microdomain at a fixed status of their extraction. Further, the use of our *in vitro* setup in which actin patches associated with synapses, would be expected to be lost during synaptosome-preparation, provides a controlled, artificial system in which all postsynaptic densities adopt the characteristics of dendritic shaft synapses, accessible to the MT cytoskeleton. In neurons, synapses may be more complex and additional factors may play a role in determining MT dynamics that may be absent in our *in vitro* assays. Thus, MT growth will need to be examined in SHANK3_L68P_ PV neurons, where the effect of CAMSAP2 depletion on MT growth at synapses of PV neurons can be investigated. The increased abundance of the nucleation factor γ-tubulin in the vicinity of PSDs of PV neurons may further play a role in the nucleation of MTs.

Interestingly, there are a handful of MT associated proteins (MAPs) implicated to play a role in ASD (*59*) and many of them are implicated in neurite outgrowth (*59*) which crucially depends on microtubule polarity. This is consistent with our observation of the overlapping interactome of CAMSAP2 and SHANK3 and extends the intersection between the postsynaptic density and the MT cytoskeleton to a potential target in ASD.

To get an idea what effect this mechanism would have in the disorder context, underlying changes in protein composition were additionally investigated in homozygous SHANK3_WT_, SHANK3_L68P_ and heterozygous samples. Comparing the results of our *in vitro* MT experiments with the quantification of selected protein levels from forebrain synaptosome samples gave us further insights on how this mutation alters the synaptic composition. While most protein levels were unchanged in total brain lysates, the trend for SHANK2 levels showed a decrease from SHANK3_WT_ to homozygous SHANK3_L68P_ samples. In synaptosomes, this trend was reversed with higher levels of SHANK2 in the SHANK3_L68P_ condition. We saw the same trend with SHANK1 and CAMSAP2 in synaptosome samples. All of these proteins interact with SHANK3 while SHANK1 and SHANK2 can redundantly fill out roles SHANK3 has (*60*). This might indicate that an upregulation of SHANK1 and SHANK2 expression, increase in localisation to the synapse or decrease of ubiquitination are made in an attempt to compensate for changes introduced by replacing SHANK3_WT_ with the SHANK3_L68P_ mutation.

Our hypothesis that the SHANK3_L68P_ and CAMSAP2 interaction plays a large role in PV neurons is supported by the observation that alterations of SHANKs in GABAergic or even PV neurons often recapitulate effects seen in pan-neuronal alterations (*61*, *62*). Traditionally, like many (excitatory) synaptic proteins, SHANK3 has been best studied in excitatory neurons. There is however mounting evidence that especially in ASD, the role of SHANKs in GABAergic neurons has been underestimated. A knockout (KO) of exons 14-16 of Shank3 was compared under a GABAergic neuron specific promotor to a global KO. In most behavioural analyses, the behavioural deficits were replicated by the GABAergic-specific KO (*61*), highlighting the importance of SHANK3 in GABAergic cells. A recent paper further highlights the importance of SHANK3 specifically in parvalbumin-positive GABAergic neurons. When a KO of exon 11 of SHANK3 was performed in mice, ASD-related behaviour and a decrease in the activity of PV neurons could be observed (*62*).

In summary, we propose a mechanism by which dendritic MTs can be tethered at the PSD of shaft synapses in PV neurons. The increase in synapse-MT association in the context of the disorder-associated SHANK3L68P mutation might alter the stability of dendritic MTs, in concert thereby leading to alterations in dendritic complexity.

## Materials and Methods

### Dissociated hippocampal neuron cultures from E18/19 and P0 rats

Coverslips heated to 160-180°C for 3 h were placed in their corresponding well plates (18 mm for 12 wp) and incubated with poly-L-Ornithine (PLO) in borate buffer (pH 8.1) followed by an incubation with Laminin. Before seeding neurons, they were washed multiple times with MilliQ water and HBSS.

P0 or embryonal day 18/19 (E18/19) Wistar (RjHan:WI) rats were decapitated and the brain and the hippocampi isolated. E18/19 rats were prior removed from a decapitated adult rat. All tissue isolation steps were done in ice-cold HBSS. Once hippocampi from all animals were collected in 10-15 ml HBSS, they were carefully washed 5x with 9 ml ice-cold HBSS under a flow-hood. For each hippocampus, 100 µl 0.25% Trypsin/EDTA solution was added to the hippocampi which were suspended in 2 ml HBSS. After 15 min at 37°C, Trypsin was removed by washing 5x with 9 ml 37°C warm HBSS. 2 ml remaining HBSS-tissue mixture were moved to a 2 ml reaction tube and triturated with 20G/26G needles. The cell solution was filtered (Easy Strainer, Greiner 542000) and the filter washed with HBSS. Cells were then manually counted in a Neubauer chamber and plated in densities between 40000-80000 cells/ml in prewarmed full medium (Dulbecco’s modified Eagle’s medium (DMEM), 10% fetal bovine serum (FBS), 1x penicillin/streptomycin, 2 mM glutamine). After 1 h at 37°C and 5% CO_2_ the medium was exchanged to 37°C warm Brain Phys medium (Gibco) with added 1x SM1 and 0.5 mM glutamine supplements. 24-48 h later, 2 µM Cytosine β-D-arabinofuranoside (AraC) was added to the cultures to hinder glia-cell proliferation. To counteract a change in osmolarity due to evaporation of the medium, after one week 10% medium was removed and 20% fresh medium added.

### Transfection of dissociated neurons

To introduce DNA plasmids, we used lipofectamine transfection. Briefly, DNA and lipofectamine 2000 were each diluted in prewarmed BrainPhys and incubated at room temperature for 5 min. After mixing DNA and lipofectamine, the solution was incubated for 20 min. The mixture was gently added to neurons in serum-free medium at DIV9 for CAMSAP2 MBD and GFP transfections for Sholl analyses. After around 90 min incubation, the medium was removed and replaced with BrainPhys +/+. Neurons were placed into the incubator until DIV15 when they were fixed for immunostaining.

### Primary Hippocampal Neuron Culture

Primary hippocampal neurons were prepared from embryonic day 18 (E18) embryos obtained from the breeding of *SHANK3_L68P_* heterozygous mice. Hippocampi were dissected in ice-cold plain Neurobasal medium (Gibco). Genotyping was performed for each embryo, and only tissues from wild-type (WT) and *SHANK3_L68P_* embryos were selected for further processing. The selected hippocampal tissues were enzymatically digested with papain (Worthington) for 20 min at 37°C and then mechanically dissociated by trituration (10–20 strokes).

The dissociated neurons were seeded onto 18-mm coverslips coated with poly-D-lysine at a density of 100000 cells per coverslip. The initial plating medium consisted of Neurobasal medium supplemented with 5% FBS, 1% B-27, 1 mM Glutamax, and 1 mM sodium pyruvate. Four hours after plating, the medium was replaced with neuron culture medium (Neurobasal medium, 1x B-27, 1x Glutamax, and 1 mM sodium pyruvate). The cultures were maintained without media exchange until DIV7.

### AAV Transduction

At DIV7, 50% of the culture medium was replaced with fresh culture medium. During this medium exchange, neurons were transduced with AAV-php.eb-S5E2-mCherry-P2A-Cre and either AAV-php.eb-hSyn-Mir30-eGFP-shCAMSAP2 (shCAMSAP2) or its corresponding scrambled control (AAV-php.eb-hSyn-Mir30-eGFP-shCAMSAP2-scrambled) at a titer of 5×10^8^ genome copies (GC) per coverslip. The neurons were subsequently fixed and processed for immunostaining at DIV15.

### Cell lines

HEK293T (from (*63*) were grown in Full medium (Dulbecco’s modified Eagle’s medium (DMEM), 10% fetal bovine serum (FBS), 1% Penicilin/Streptavidin). Newly thawed cells were grown in 20% until split for the second time. To split cells, old medium was removed from cells, and they were carefully washed with 37°C warm HBSS. 2 ml 0.05% 37°C warm Trypsin/EDTA were added to the cells and after a few minutes 8 ml Full medium were pipetted on the cells, and they were separated from the plastic surface. From this cell homogenate, new HEK293T cell dishes were prepared in the desired density. They were grown at 37°C 5% CO_2_ until usage.

### Cell line transfections

Cells were transfected with MaxPEI (MW 40 kDa, pH) according to the manufacturer’s guidelines in a DMEM/Ham’s F10 medium including FBS and Penicilin/Streptomycin. For proteins intended to be affinity-purified with Streptavidin-beads a biotin ligase (BirA) was coexpressed with 3 µg/10 cm dish. The protein of interest was transfected with 10-20 µg plasmid DNA/10 cm dish (Table 4).

### Immunoblotting

Samples were run on self-cast 8 – 12% SDS acrylamide gels or precast gradient gels (BioRad) depending on the weight of the protein of interest. 2,2,2-Trichlorethanol was added to the gels to allow for detection of total protein with the help of UV-light. Samples were prepared by denaturing after the addition of 2x or 4x SDS-loading buffer at 98°C for 5-10 min. Throughout the stacking gel, proteins were run at 80 V which was increased up to 120 V for the running gel.

From SDS-acrylamide gels, samples were blotted onto low fluorescence or conventional PVDF membranes for fluorescence and chemiluminescent sample detection, respectively. For wet tank immunoblotting, the western blot sandwich was assembled in in ice cold blotting buffer (192 mM Glycine, 0.1% (w/v) SDS, 15% (v/v) methanol, 25 mM Tris-base, pH 8.3).

For semi-dry immunoblotting using the Trans-Blot® Turbo™ 1x Transfer System Trans-Blot Turbo Buffer (BioRad) was used with supplemented ethanol (40%) (Turbo buffer). To prepare PVDF membranes (PVDF-LF when used for fluorescent detection of samples), membranes were first incubated with methanol for 40 s, 2 min with MilliQ water and 5 min with Turbo buffer. To assemble the Western sandwich, two Whatman papers soaked in Turbo buffer were laid on each electrode. The membrane was gently laid on the Whatman paper on the cathode (+) and the ready SDS acrylamide gel laid on top. The other two Whatman papers were placed on top and air bubbles removed gently with pressure. The chamber was closed, and the appropriate transfer program was selected.

After blocking membranes in 5% skim milk in Tris-buffered saline with Tween (TBS-T; 20 mM Tris pH 7.4, 150 mM NaCl, 0.1% Tween-20) or EveryBlot blocking buffer (12010020, BioRad), membranes were incubated with primary antibodies (diluted according to Table 1) in TBS-A (TBS pH 7.4, 0.02% sodium-azide) under constant rotation overnight at 4°C. After several TBS and TBS-T washes, suitable secondary antibodies conjugated to horseradish peroxidase (HRP) or fluorescent dyes (Table 2) were applied for min 60 min at room temperature (RT) in 5% skim milk in TBS-T. After three washes with TBS and TBS-T, the membranes were developed after the addition of ECL (100 mM Tris-HCl pH 8.5), 5.3 mM H_2_O_2_, 1.25 mM luminol, 2 mM 4-Iodophenylboronic acid (4-IPBA)) using a ChemoCam imager (Intas) or ChemiDoc MP system (BioRad).

**Table 1:** Primary Antibodies.

| Antibody | Species | Application and Dilution | Source | Identifier |
| --- | --- | --- | --- | --- |
| CAMSAP2 | rabbit | coIP<br>WB 1:2000<br>ICC 1:100 | NovusBio | NBP1-21402 |
| CAMSAP1 | rabbit | coIP<br>WB 1:5000 | NovusBio | NBP1-26645 |
| CAMSAP3 | rabbit | coIP<br>WB 1:1000 | Biozol (Abcepta) | AP18323a-ev |
| SHANK3 | guinea pig | ICC 1:500 | SySy | 162304 |
| SHANK2 | rabbit | WB 1:500 | SySy | 162202 |
| SHANK1 | rabbit | WB 1:500 | SySy | 162002 |
| Parvalbumin | chicken | ICC 1:500 | NovusBio | NBP2-50036 |
| Parvalbumin | guinea pig | ICC 1:500 | SySy | 195004 |
| Homer 1/2/3 | rabbit | ICC 1:500 | SySy | 160103 |
| PSD95 | mouse | ICC 1:500 | NeuroMab | 75-028 |
| vGlut1 | guinea pig | ICC 1:500 | NanoTag, SySy | N1602-SC3 |
| Calbindin-D28k | guinea pig | ICC 1:500 –<br>1:300 | SySy | 214004 |
| Calretinin | mouse | ICC 1:500 | SySy | 214111 |
| MAP2 | rabbit | ICC 1:500 | SySy | 188002 |
| MAP2 488 | mouse | ICC 1:500<br>IHC 1:500 | Merck Millipore | MAB3418X |
| GFP | mouse | WB 1:5000 | Biolegends | 902601 |

**Table 2:** Secondary Antibodies.

| Antibody | Conjugate Fluorophor | Species | Application and Dilution | Source | Identifier |
| --- | --- | --- | --- | --- | --- |
| Anti-mouse | Alexa Fluor 405 | Goat | ICC 1:500<br>IHC 1: 500 | Abcam | ab175660 |
| Anti-mouse | Alexa Fluor 488 | Goat | ICC 1:500<br>IHC 1: 500 | Life Technologies | A-11001 / A11029 |
| Anti-mouse | Alexa Fluor 568 | Goat | ICC 1:500<br>IHC 1: 500 | Life Technologies | A-11004 / A11031 |
| Anti-mouse | Abberior STAR 580 | Goat | ICC 1:500<br>IHC 1: 500 | Abberior | ST580-1001-500µg |
| Anti-rabbit | Alexa Fluor 405 | Goat | ICC 1:500<br>IHC 1: 500 | Abcam | ab175652 |
| Anti-rabbit | Atto 647N | Goat | ICC 1:500<br>IHC 1:500 | Sigma | 40839 |
| Anti guinea pig | Alexa Fluor 405 | Goat | ICC 1:500 | Abcam | ab175678 |
| Anti-guinea pig | Alexa Fluor 488 | Goat | ICC 1:500 | Invitrogen | A-11073 |
| Immunoblotting |  |  |  |  |  |
| Anti-mouse igG | Star Bright Blue 700 | Goat | WB 1:5000 | BioRad | 12004159 |
| Anti-mouse igG | Star Bright Blue 520 | Goat | WB 1:5000 | BioRad | 12005866 |
| Anti-rabbit igG | Star Bright Blue 700 | Goat | WB 1:5000 | BioRad | 12004161 |
| Anti-rabbit igG | Star Bright Blue 520 | Goat | WB 1:5000 | BioRad | 12005869 |
| Anti-guinea pig | IRDye® 800CW | Donkey | WB 1:5000 | LiCor | 926-32411 |
| Anti-guinea pig | DyLight 800 | Goat | WB 1:5000 | Invitrogen | SA5-10100 |
| Anti-guinea pig | DyLight 650 | Goat | WB 1:5000 | Invitrogen | SA5-10097 |
| Anti-guinea pig | HRP | Donkey | WB 1:10000 | Dianova | 706-035-148 |
| Anti-mouse | HRP | Goat | WB 1:10000 | Dianova | 115-035-146 |
| Anti-rabbit | HRP | Goat | WB 1:10000 | Dianova | 111-035-144 |
| Anti-goat | HRP | Donkey | WB 1:10000 | Dianova | 705-035-147 |

### Brain lysate preparation

To obtain brain lysate to be used in co-immunoprecipitations (coIPs), the brain of a pregnant rat was isolated after CO_2_/O_2_ sedation and decapitation. Bulbus olfactorius and cerebellum were removed. 9 ml tissue lysis buffer (20 mM Tris-HCl pH 7.4-8.0, 150 mM NaCl, 1% TritonX-100, 1x protease inhibitors without EDTA) was added per 1 g tissue. The solution was homogenised using a Dounce homogeniser. To remove debris, the sample was spun down at 900 rcf at 4°C for 15 min. Aliquots of 1-2 ml were prepared and either shock-frozen in liquid nitrogen to be stored at −80°C or used immediately.

### Synaptosome preparation/Subcellular fractionation

To evaluate protein levels in synapses, synaptosomes were isolated using subcellular fractionation (***S***5A). First, brains of 3-month-old mice were isolated and bulbus olfactorius and cerebellum removed. The remaining tissue was homogenised in 10 ml/g buffer A (0.32 M sucrose, 5 mM HEPES, pH 7.4) and centrifuged at 1000 rcf for 10 min. The pellet was resuspended and re-centrifuged. The pellet containing the nuclei was discarded, and supernatants combined. To obtain a pellet containing the heavy membrane fraction (P2), the supernatants were centrifuged at 12000 rcf for 20 min. The pellet was rehomogenised in buffer B (0.32 M sucrose, 5 mM Tris, pH 8.1) and carefully loaded on a sucrose gradient from 0.85 M, 1 M to 1.2 M. After spinning the sample at 85000 rcf for 2 h, the phase between the 1 M and 1.2 M sucrose that contains the synaptosome fraction was collected. Protein concentrations were analysed with Amido Black and samples treated with SDS LB prior for further analysis using immunoblotting.

### Protein concentration measurement

Protein concentration was determined with the help of a standard curve using differently concentrated BSA-samples. Amidoblack was added to the samples. After 20 min the samples were centrifuged at 16000 rcf for 5 min. Pellets were subsequently washed three times with a 10% acetic acid 90% methanol solution. The pellet was then airdried and solved in 0.1 M NaOH. 30 min later the OD was measured at 620 nm and a standard curve calculated.

### Co-Immunoprecipitations (coIPs)

#### Bead preparation

Unless described otherwise, all steps were performed on ice or at 4°C.

For endogenous co-immunoprecipitations an equal mixture of magnetic protein A&G beads were used. For the purification of biotinylated proteins, Dynabeads (Streptavidin-beads) were used instead.

First, beads were washed 3x in washing buffer in a 2 ml reaction tube (washing buffer: 20 mM Tric-HCl pH 7.4-8.0, 150 mM NaCl, 0.1% TritonX-100). To avoid unspecific binding the beads were blocked in blocking buffer (20 mM Tris-HCl pH 7.4-7.8, 150 mM NaCl, 0.2 µg/µl chicken egg albumin (CEA)) for at least 1 h under constant rotation. The beads were then washed 3x in washing buffer. A defined volume of supernatant was kept on the beads to allow for precise distribution to each individual sample.

#### Endogenous co-immunoprecipitation (endo coIP)

Unless described otherwise, all steps were performed on ice or at 4°C. To allow the primary antibody raised against the protein of interest to be bound to its target, adult rat brain lysate or adult mouse SHANK3_L68P_ KI (WT/het/hom) brain lysate and 2-3 µg of antibody were incubated under constant rotation for 2 h. Then the beads were added to the antibody-brain lysate mixture and incubated under constant rotation for 3-4 h. The beads were then washed five times with washing buffer. The washed beads were resuspended in 40 µl preheated SDS loading buffer and denatured at 95°C for 10 min.

#### Heterologous coIPs

To harvest proteins purified from transfected HEK cells, they were washed with ice-cold TBS. Cells on 10 cm dishes were lysed with 400 µl HEK lysis buffer containing protease inhibitors (EDTA-free) and scraped off. To remove cell debris, the lysate was spun down at 13000 rcf for 15 min at 4°C. The lysate was then incubated with the blocked beads and incubated overnight under constant rotation. Both bait protein and interactor were pulled down from HEK cell lysate. The beads were then washed 5x with HEK lysis buffer. 40 µl SDS loading buffer were added to the beads after removing the last wash step and denatured at 98°C for 10 min.

To find out the SHANK3 binding site on CAMSAP2, CAMSAP2 truncations were co-expressed with a biotin ligase in HEK293T cells as described before. CoIP was performed as described before, but instead the interacting protein SHANK3 was bound to CAMSAP2 truncations by adding brain lysate to the beads for 1 h under constant rotation. Previously, CAMSAP2 truncations were bound to the Streptavidin beads for 2 h under constant rotation.

#### Heterologous coIPs (SHANK3 WT vs L68P binding to CAMSAP2)

A difference in the binding to CAMSAP2 between WT SHANK3 and SHANK3 harbouring the L68P mutation was investigated by expressing SHANK3 1-676 AA constructs (WT or L68P) in HEK cells, similarly as described in “Overexpression coIPs” (SHANK3 binding site on CAMSAP2). Brain lysate was added to the beads o/n under constant rotation.

### Protein purification from *E. coli* – SHANK3 1-676 WT and L68P

To obtain larger quantities of recombinant proteins, *E.coli* BL21 bacteria carrying plasmids enabling IPTG-inducible protein expression were used. Cultures were inoculated with a pre-culture. Upon reaching an OD around 0.6, they were induced with 1 mM IPTG. They were grown in LB medium with the antibiotic they had been selected for at 18°C o/n. The bacterial pellets were spun down at 13000 rcf for 15 min at 4°C and stored at −20°C until further usage. Pellets were thawn on ice and resuspended with PBS resuspension buffer. To lyse the samples, lysozyme was added. After 30 min incubation on ice, the lysates were frozen at −80°C at least o/n. Upon continuation, lysates were thawn on ice. Benzonase (c_final_ = 50 U/ml) were added to the thawn lysate and incubated for at least 1 h on ice to digest genomic DNA and reduce the viscosity of the sample. To remove insoluble cell-debris, the lysate was spun down at 11000 rcf for 1 h at 4°C and the supernatant transferred to a pre-chilled tube. To allow the His-tagged protein to be purified, a Ni-NTA column was prepared in parallel. 3 ml of the bead-mixture were mixed with ∼10 ml PBS resuspension buffer and the column allowed to empty under gravity flow after the beads had settled. The bacteria lysate was allowed to incubate with the beads for 10 min, before being allowed to enter by gravity flow. To remove unspecifically bound proteins, the column was washed multiple times with PBS washing buffer (+) and PBS washing buffer (-). Elution was performed using PBS elution buffer containing SenP2 protease cleaving the SUMO-tag located between the protein of interest and the His-tag bound to the beads.

### Protein purification from HEK293T cells

Beads were blocked as described before. HEK293T cells were transfected as described previously. After 12-36 h (depending on the protein), cells were harvested. All following steps were performed on ice. First, medium was removed and cells washed with ice-cold HBSS. Cells were then lysed in HEK293T lysis buffer and detached from the surface. The lysate was collected in a 1.5 ml tube and centrifuged at 13000 rcf for 15 min to remove cell debris. 40 µl supernatant were collected for an SDS gel/immunoblot analysis. Blocked and washed beads were distributed equally between samples and incubated with supernatant from the previous centrifugation step for 1 h under constant rotation. The beads were then washed five times with HEK293T lysis buffer (without protease inhibitors) and two times with TEV protease buffer. Proteins were then eluted in 50 µl TEV protease buffer containing 2 µg TEV protease while shaking. Proteins were harvested and beads removed after 18-36 h.

### *In vitro* MT co-pelleting assay

For the *in vitro* MT co-pelleting assay, first microtubules were synthesised by mixing bovine tubulin, G-PEM, GMPCPP (a stable GTP derivate, Table 3) and PEM80 (Table 5).The mixture was incubated at 37°C for 20 min and kept at room temperature afterwards to avoid MT depolymerisation.

**Table 3:** Drugs/Agents/Chemicals.

| Drug/Enzymes etc. | Dilution | Source | Identifier |
| --- | --- | --- | --- |
| cOmplete(TM), EDTA-free Protease Inhibitor | 1/ 50 ml | Sigma (Roche) | 4693132001 |
| Benzonase |  | Santa Cruz | SC-202391 |
| TEV protease | 2 µg/50 µl | Sigma | T4455-1KU |
| MaxPEI (40 kDa) |  | Polysciences | 24765/ 23966-100 |
| SureBeads™ Protein A<br>Magnetic Beads |  | BioRad/Invitrogen | 1614013/10001D |
| SureBeads™ Protein G<br>Magnetic Beads |  | BioRad/Invitrogen | 1614023/10003D |
| Streptavidin 280M beads |  | ThermoFisher (Invitrogen) | 11206D |
| GFP trap beads |  | Chromotek | Gtma-20 |
| Chicken egg white (CEA) | Stock<br>concentration<br>2 µg/µl | Sigma | A7641-50MG |
| Bovine serum albumin (BSA) |  | Roth | 8076.2 |
| Poly-L-Lysine (PLL) |  | Sigma | P2636-25mg |
| Poly-D-Lysine (PDL) | 10mg/ml (100X) | Sigma | P7889-100mg |
| porcine brain tubulin protein |  | Cytoskeleton/Tebu Bio | T240-B |
| Lysozyme | 20 mg/ 10 ml | Sigma | 62970/L6876 |
| Lipofectamine 2000 |  | ThermoFisher | 11668019 |
| Glycerol |  | Roth | 3783,1 |
| EDTA |  | Roth | 8040.3 |
| EGTA |  | VWR | 0732-100g |
| Kynurenic acid |  | biomol (Cayman<br>chemicals) | Cay16792-500 |
| Sucrose (Saccharose D-(+)) |  | VWR | 27480294 |
| Glucose (Glucose D-(+)) |  | Roth | X997.2 |
| Roti Histofix 10% (PFA) |  | Roth | A146.6 |
| GmPcPP |  | Jena-Bio | NU-405L |
| Amidoblack powder |  | Merck microscopy | 1.01167.0025 |
| Mowiol |  | Roth | 713,2 |
| DAPI |  | Sigma | D9542-10mg |
| Laminin |  | Bio-Techn<br>Corning | 3400-010-02 1mg<br>CLS354232 |
| PLO (poly-l-ornithine) |  | Sigma | P3655-50mg |

**Table 4:** Plasmids.

| **Name** | **Promotor** | **Description/Purpose** |
| --- | --- | --- |
| Biotinylase A (BirA) | CMV | Helper construct for affinity purification |
| eGFP-BioTag | CMV | Eukaryotic eGFP expression |
| mRuby3-GCN4_P1-MacF18-DTE | hSynapsin | Live imaging of neuronal microtubules |
| His6-SUMO-SHANK3(1-676, wt) | T7 | Bacterial expression of SHANK3 1-676 |
| AviTag-GFP-SHANK3(1-676) WT | CMV | Eukaryotic expression of SHANK3 1-676 |
| His6-SenP2 | T7 | Bacterial expression of SenP2 for purifications |
| CAMSAP2 FL human | CMV | Eukaryotic CAMSAP2 FL expression |
| CAMSAP2 1-1286 human | CMV | Eukaryotic CAMSAP2 1-1286 expression |
| CAMSAP2 477-1489 human | CMV | Eukaryotic CAMSAP2 477-1489 expression |
| CAMSAP2 922-1489 human | CMV | Eukaryotic CAMSAP2 922-1489 expression |
| CAMSAP2 1349-1489 human | CMV | Eukaryotic CAMSAP2 1349-1489 expression |
| CAMSAP2 922-1034 human (= Microtubule binding domain) | CMV | Eukaryotic CAMSAP2 922-1034 expression |
| CAMSAP2 1-476 human | CMV | Mammalian CAMSAP2 1-476 expression |
| mRuby3-MACF18 | EF1a | Microtubule live imaging in floxed cells. |
| AviTag-GFP-SHANK3(1-348) WT | CMV | Eukaryotic expression of SHANK3 1-348 |
| AviTag-GFP-SHANK3(349-676) WT | CMV | Eukaryotic expression of SHANK3 349-676 |
| CAMSAP2 922-1034 human (= Microtubule binding domain)<br>mRuby3 | CMV | Eukaryotic CAMSAP2 922-1034 expression |
| CAMSAP2 922-1034 human (= Microtubule binding domain)<br>tagBFP | CMV | Eukaryotic CAMSAP2 922-1034 expression |
| AviTag-GFP-SHANK3(SPN domain) WT | CMV | Eukaryotic expression of SHANK3 SPN domain |
| AviTag-GFP-SHANK3(ARR) WT | CMV | Eukaryotic expression of SHANK3 ARR |
| CAMSAP2 922-1034 human (= Microtubule binding domain) mEmerald | CMV | Eukaryotic CAMSAP2 922-1034 expression |
| CAMSAP2 FL human - mRuby3 | CMV | Eukaryotic CAMSAP2 FL expression |
| pEGFP-C2-AviTag-GFP-TEV-HA-SHANK3 (1-676) WT | CMV | Eukaryotic SHANK3 1-676 expression |
| pEGFP-C2-AviTag-GFP-TEV-HA-SHANK3 (1-676) R12C | CMV | Eukaryotic SHANK3 1-676 expression |
| CAMSAP2 FL human - mRuby3 | fPV | Eukaryotic CAMSAP2 FL expression |
| (ccr5tc.fPV)-PSD95fingR(GFP)-IRES.MacF18(mRuby3) | fPV | Endogenous PSD95 labelling and microtubule live imaging in Parvalbumin neurons |
| CAMSAP2 922-1034 human (= Microtubule binding domain) human - GFP | fPV | Eukaryotic CAMSAP2 922-1034 expression |
| CAMSAP2 922-1034 human (= Microtubule binding domain) human - mRuby3 | fPV | Eukaryotic CAMSAP2 922-1034 expression |
| CAMSAP2 922-1034 human (= Microtubule binding domain) human - tagBFP | fPV | Eukaryotic CAMSAP2 922-1034 expression |
| CAMSAP2 922-1034 human (= Microtubule binding domain) human - mEmerald | fPV | Eukaryotic CAMSAP2 922-1034 expression |

**Table 5:**
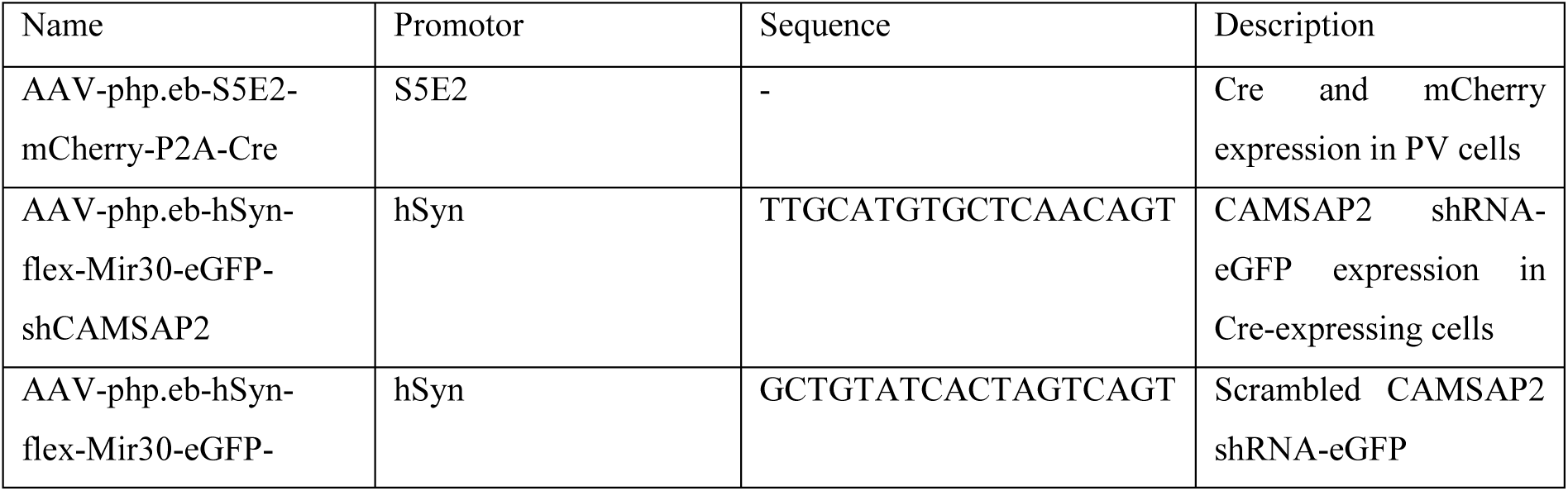
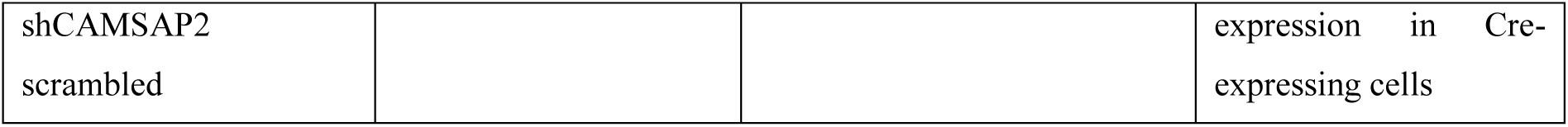
Viruses.

All proteins that would be used in the assay were centrifuged for 100 000 rcf for 5 min at 4°C. To have proteins of interest bind to the microtubules (directly or indirectly), microtubules were incubated with purified proteins in PEM80 buffer. For each protein mixed with microtubules, the same mixture but without microtubules serves as a negative control. The protein-microtubule mixtures were incubated for 30 min at room temperature. They were then gently pipetted onto cushion buffer (60% glycerol in PEM80). The samples were then centrifuged at room temperature (21°C) for 40 min at 100 000 rcf (Beckmann TLA Ultracentrifuge TD503 rotor TLA100.3). Of each sample 50 µl were taken off to keep as supernatant in which proteins that did not bind to microtubules should be found if MTs were present. The middle 50 µl were discarded and the last 50 µl were kept as the pellet sample. All samples were then denatured at 98°C for 10 min with 2x SDS loading buffer. Acquired samples were then analysed on gels and western blots.

### Mass spectrometry – SHANK3 1-676 pulldown

For the Bio-GFP-SHANK3 interaction and phosphorylation state analysis, SDS-PAGE gel lanes were cut into 2 mm slices and subjected to in-gel reduction with dithiothreitol, alkylation with iodoacetamide and digested with trypsin (sequencing grade; Promega). Nanoflow liquid chromatography tandem mass spectrometry (nLC-MS/MS) was performed on an EASY-nLC coupled to an Orbitrap Fusion Tribrid mass spectrometer (ThermoFisher Scientific) operating in positive mode. Peptides were separated on a ReproSil-C18 reversed-phase column (Dr Maisch; 15 cm × 50 μm) using a linear gradient of 0–80% acetonitrile (in 0.1% formic acid) during 90 min at a rate of 200 nl/min. The elution was directly sprayed into the electrospray ionisation (ESI) source of the mass spectrometer. Spectra were acquired in continuum mode; fragmentation of the peptides was performed in data-dependent mode by HCD.

For global quantitative proteomics analysis, proteins in whole cell extracts (WCE) from pyramidal neurons were first reduced with 5 mM DTT and cysteine residues were alkylated with 10 mM iodoacetamide. Protein pellets were then dissolved in 1 ml 50 mM Tris/HCl pH 8.2, 0.5% sodium deoxycholate (SDC) and proteins were digested with LysC (1:200 enzyme:protein ratio) for 4 h at 37°C. Trypsin was added (1:100 enzyme:protein ratio) and the digestion proceeded overnight at 30°C. Digests were acidified with 50 μl 10% formic acid (FA) and centrifuged at 8,000 rcf for 10 min at 4 °C to remove the precipitated SDC. The supernatant was transferred to a new centrifuge tube. The digests were purified with C18 solid phase extraction (Sep-Pak, Waters), lyophilised and stored at −20°C.

Proteolytic peptides were then labelled with isobaric TMT 10-plex labelling reagents (Thermo Scientific) allowing for peptide quantitation. Peptides were mixed at the 10-plex level and further fractionated by HILIC chromatography into six fractions. Fractions were collected and analysed by nanoflow LC-MS/MS. nLC-MS/MS was performed on EASY-nLC 1200 coupled to an Orbitrap Lumos Tribid mass spectrometer (Thermo) operating in positive mode and equipped with a nanospray source. Peptides were separated on a ReproSil C18 reversed phase column (Dr Maisch GmbH; column dimensions 15 cm × 50 µm, packed in-house) using a linear gradient from 0 to 80% B (A = 0.1% formic acid; B = 80% (v/v) acetonitrile, 0.1% formic acid) in 70 min and at a constant flow rate of 200 nl/min using a splitter. The column eluent was directly sprayed into the ESI source of the mass spectrometer. Mass spectra were acquired in continuum mode; fragmentation of the peptides was performed in data-dependent mode using the multinotch SPS MS3 reporter ion-based quantification method with HCD activation type.

### Mass spectrometry analysis and statistics

Raw mass spectrometry data were analysed with the MaxQuant software suite as described previously with the additional options ‘LFQ’ and ‘iBAQ’ selected. The A false discovery rate of 0.01 for proteins and peptides and a minimum peptide length of 7 amino acids were set. The Andromeda search engine was used to search the MS/MS spectra against a Uniprot database, taxonomy Mus musculus (version January 2019) concatenated with the reversed versions of all sequences. A maximum of two missed cleavages was allowed. The peptide tolerance was set to 10 ppm and the fragment ion tolerance was set to 0.6 Da for HCD spectra. The enzyme specificity was set to trypsin and cysteine carbamidomethylation was set as a fixed modification, while methionine oxidation and phosphorylation (STY) were set as variable modifications. Both the PSM and protein FDR were set to 0.01. In case the identified peptides of two proteins were the same or the identified peptides of one protein included all peptides of another protein, these proteins were combined by MaxQuant and reported as one protein group. Known contaminants and reverse hits were removed. Both the ‘evidence’ and ‘modificationsspecificpeptides’ output text files were used for further in-depth analysis.

For quantitative proteomic analysis, data were processed with Proteome Discoverer 2.3 (ThermoFisher Scientific, Table 6). The Mascot search algorithm (version 2.3.2; MatrixScience) was used for searching against a Uniprot database. The peptide tolerance was set to 10 ppm and the fragment ion tolerance was set to 0.8 Da. A maximum number of 2 missed cleavages by trypsin were allowed and carbamidomethylated cysteine and oxidised methionine and phosphorylation (STY) were set as fixed and variable modifications, respectively. A target / decoy database search strategy was applied with a strict FDR of 0.01 and a relaxed FDR of 0.05 and Percolator validation of PSMs. Typical contaminants were omitted from the output tables.

**Table 6:** Buffers and media.

| Buffer | Used for | Ingredients |
| --- | --- | --- |
| Tissue lysis buffer | Endo coIP | 20 mM Tris-HCl pH 7.4-8.0, 150 mM NaCl, 1% TritonX-100, 1x protease inhibitors without EDTA |
| Washing buffer | Endo coIP, coIP | 20 mM Tric-HCl pH 7.4-8.0, 150 mM NaCl, 0.1% TritonX-100 |
| Blocking buffer | Endo coIP, coIP | 20 mM Tric-HCl pH 7.4-8.0, 150 mM NaCl, 0.2 µg/µl CEW |
| HEK lysis buffer | coIP | 50 mM HEPES, 300 mM NaCl, 0.5% Triton X-100, pH 7.4, 1x protease inhibitor without EDTA |
| TEV protease buffer | Protein purification | 50 mM HEPES, 150 mM NaCl, 1 mM MgCl<sub>2</sub>, 1 mM EGTA, 0.05% Triton X-100, 1 mM DTT, pH 7.4 |
| Blocking buffer (BB-HE) | ICC, IHC | 10% heat inactivated horse serum, 0.1% Triton X-100 in PBS |
| DMEM +/+ (Full medium) | Cell line cultivation, plating of dissociated neurons | 10% FCS (fetal calf serum), 1x Pen/Strep in DMEM |
| BrainPhys | Transfection of dissociated neurons | StemCell 5792 |
| BrainPhys +/+ | Cultivation of dissociated neurons | Brain Phys (StemCell 5792) supplemented with 1x SM1 and 0.5 mM glutamine |
| Neurobasal medium +/+ | Cultivation of dissociated mouse neurons | Neurobasal medium (Gibco 21103049) supplemented with 1x B-27, 1x GlutaMax (Gibco 35050061) and 1 mM Sodium pyruvate (Gibco, 11360070) |
| Ham's F10 Nutrient Mix |  | Gibco™ 11-550-043 |
| PEM80/MRB80 | Microtubule co-sedimentation assay | 80 mM PIPES, 1 mM EGTA, 4 mM MgCl<sub>2</sub>, pH 6.8 in ddH<sub>2</sub>O |
| G-PEM | Microtubule co-sedimentation assay | 10% glycerol in PEM80 |
| Cushion buffer | Microtubule co-sedimentation assay | 60% glycerol in PEM80 |
| 4x SDS sample buffer | Denaturing of peptides for SDS-PAGE | 1% (w/v) SDS, 40% (v/v) Glycerol, 20% (v/v) β-mercaptoethanol, 0,004% (v/v) Bromophenol Blue |
| SDS-PAGE running buffer | SDS-PAGE | 192 mM glycine, 0.1% (w/v) SDS, 25 mM Tris-base, pH 8.3 |
| Blotting buffer (WB) | Immunoblotting | 192 mM Glycine, 0.2% (w/v) SDS, 20% (v/v) Methanol, 25 mM Tris-base, pH 8.3 in dH<sub>2</sub>O |
| Blocking buffer (WB) | Reducing unspecific band detection | 5% milk (w/v) in TBS-T |
| Blotting buffer (semi-dry) |  | 1x TransBlot Turbo, 40% EtOH ad. 1000 ml dH<sub>2</sub>O |
| TBS | PVDF membrane washes |  |
| TBS-T | PVDF membrane washes | 1% Tween in TBS |
| TBS-A | Primary antibody aliquot storage for immunoblotting | 0.1% sodium-azide in TBS |
| Buffer A | Subcellular fractionation | 0.32 M sucrose, 5 mM HEPES, pH 7.4<br>1x protease inhibitor |
| Buffer B | Subcellular fractionation | 0.32 M sucrose, 5 mM Tris/HCl, pH 8.1 |
| Master buffer | *In vitro* microtubule growth assay | 75 mM KCl, 1 mM GTP, 0.2% Methylcellulose, 0.5 mg/ml κ-Casein, 200 μg/ml Glucose oxidase, 4 mM DTT in MRB80 |
| Final mix | *In vitro* microtubule growth assay | 30 μM tubulin, 1.2 μM HyLite647 tubulin, 40 μg/ml Catalase, 6 mg/ml Glucose |

**Table 7:** Software.

| Software | Purpose | Source |
| --- | --- | --- |
| FIJI (FIJI is just ImageJ)<br>Plugins used:<br>BioFormats<br>ComDet<br>KymoClear<br>NeuronJ | Image processing | https://imagej.net/software/fiji |
| Simple Neurite Tracer |  |  |
| Adobe Illustrator | Image processing and Figure preparation | https://www.adobe.com/de/illustrator |
| OpenView Version 1.5 | Image processing |  |
| Graphpad Prism | Statistics | https://www.graphpad.com |
| Proteome Discoverer 2.3 | Mass spectrometry analysis | ThermoFisher Scientific |

Relative peptide quantitation was based on SPS MS3 reporter ion spectral intensities within the Proteome Discoverer 2.3 framework. Both unique and razor peptides were included for quantitation, while the co-isolation threshold was set to 30% and the SPS matches threshold to 65%. Normalisation was performed on total peptide amount and protein ratios were calculated based on protein abundances.

### Mass spectrometry of CAMSAP2 pulldown

Samples used in mass spectrometry were prepared as described in Overexpression coIPs (CAMSAP2 binding site on SHANK3). They were then run on precast gels and stained with Coomassie solution overnight. After multiple washes with distilled water, in-gel digest of each lane was performed.

Mass spectra were acquired using an LTQ Orbitrap mass spectrometer and analysed with MaxQuant. The “proteinGroups.txt” output file of MaxQuant were used for further analysis.

### Immunocytochemistry (ICC) of dissociated cultures

Neurons were fixed with the addition of 4% PFA and 4% sucrose in PBS for 10 min which was subsequently washed off three times with phosphate-buffered saline (PBS). For selected antibodies, an antigen-retrieval step was included after washing which meant heating the fixed cells to 80°C for 30 min with NH_4_Cl. Alternatively, cells were fixed by adding −20°C cold methanol with 1 mM EGTA. Before proceeding with immunocytochemistry (ICC), the cells were washed multiple times with PBS for 10 min each at −20°C.

To image the MT cytoskeleton in STED resolution, soluble tubulin was removed with a brief 1 min pre-extraction using the PEM80 buffer (plus 0.3% Triton X-100, 0,1% glutaraldehyde) in at 37°C, followed by a fixation using 4% PFA, 4% sucrose in PEM80. Cells were then washed 3x with PBS for 10 min.

If not described otherwise, neurons at ∼DIV13-16 were fixed by replacing the culture medium with 4% PFA/4% sucrose. The fixative was then washed off 3x for 10 min with PBS. To permeabilise the cells, they were incubated with PBS containing 0.25% Triton X-100 which was removed with two five min wash steps with PBS. Coverslips were blocked with blocking buffer (PBS containing 10% heat inactivated horse serum (30 min at 65°C) and 0.1% Triton X-100). Coverslips were then incubated over night at 4°C with primary antibody diluted in blocking buffer (BB-HE). If used, the cells were incubated with Phalloidin at the same time. On the next day, the primary antibody was washed off with three 10 min PBS washing steps. Suitable secondary antibodies were diluted in blocking buffer and incubated for 1 h at room temperature. Secondary antibody was washed off three times for 10 min with PBS. When necessary, prelabelled primary antibodies were diluted in blocking buffer and added to the cells at this stage and incubated at 4°C over night. The antibody was then washed off with three 10 min PBS wash steps. Coverslips were then mounted on glass coverslips with the cells facing a mowiol droplet on the glass coverslip. Samples could be imaged ∼ three hours after airdrying them in the dark.

### Microscopy - Imaging

#### Spinning disk confocal imaging

Total internal reflection fluorescence (TIRF), widefield and spinning-disc confocal microscopy was performed using a custom-built inverted microscope equipped with a Nikon Eclipse Ti-E body. For widefield, an LED lamp was used. For TIRF and spinning-disc confocal microscopy, excitation lasers used were 405 nm, 488 nm, 561 nm and 640 nm.

Wide-field and spinning-disc confocal microscopy was performed with a Nikon Eclipse Ti-E controlled by VisiView software. For live imaging, samples were kept in focus with the built-in Nikon perfect focus system. The system was equipped with a 60× (Nikon, P-Apo DM 60×/1.40 oil) and a 100x TIRF objective (Nikon, ApoTIRF 100×/1.49 oil), and 488 nm, 561 nm, and 639 nm excitation lasers. Lasers were coupled to a CSU-X1 spinning disk unit via a single-mode fiber. Emission was collected through a quad band filter (Chroma, ZET 405/488/561/647 m) followed by a motorised filter wheel (Prior Scientific) with filters for CFP (480/40 m), GFP (525/50 m), YFP (535/30 m), RFP (609/54 m), and Cy5 (700/75 m, Chroma) and captured on an Orca flash 4.0LT CMOS camera (Hamamatsu). COS7 cells were imaged with the 60× objective, neurons were imaged with a 100× objective. Time-lapse images were acquired sequentially with specified intervals.

For live imaging, cells were transferred to a chamber with 5% CO_2_, 95% humidity atmosphere at 37°C. Total internal reflection fluorescence (TIRF) microscopy was performed with a 100x objective (Nikon, ApoTIRF 100×/1.49 oil).

For Sholl analyses, fixed confocal imaging was performed using a custom-built inverted microscope equipped with an Olympus IX83 body. To find transfected or transduced cells, an LED lamp was used widefield. Excitation lasers used were 405 nm, 488 nm, 561 nm and 640 nm. Lasers were coupled to a CSU-W1 spinning disk unit. Corresponding filters were used and two cameras (aCMOS pco.edge and iXON Ultra 21 Andor) were used for acquisition. Fixed imaging was performed using a 20x and a 40x objective.

#### In vitro microtubule synaptosome association assay

To investigate how synapses carrying the SHANK3 L68P mutation interact with the MT cytoskeleton, a series of washes was performed in custom-made flow channels. First, glass slides and rectangular glass coverslips were cleaned in a plasma cleaner (Harrick Plasma) for 15 min. Three flowchambers were constructed on the cleaned glass slides placing stripes of Parafilm vertically across the slides, generating a few mm long flow chambers with each one not broader than 1-2 mm resulting in a channel volume of around 10 μl. A clean glass coverslip was places on top of the Parafilm array. After gently pressing glass and parafilm against each other, the construction was placed on a 50°C warm metal plate. After 30 min the Parafilm attached the slide to the coverslip, creating flow chambers between the glass. Flow chambers were coated with 0.1 μg/μl PLL for 15 min. Following this step, all washes of flow chambers were performed by adding one channel volume of liquid on one end of the channel while simultaneously removing one channel volume from the other end of the channel. Prepared forebrain synaptosomes of SHANK3 L68P and WT mice were centrifuged for 5 min at 3000 rcf to remove aggregates prior to being added to coated flow channels. One flow channel not receiving synaptosomes served as a negative control. Synaptosomes were allowed to settle to the coated surface for 5 min. After blocking the channels with κ-Casein and a 15 min incubation, flow channels were washed 1x with MRB80. A Cy3-prelabelled PSD95-nanobody was used to label the synaptosomes for 15 min and washed 2x with MRB80 after. Tubulin-containing “final mix” containing HyLite-tubulin was added and the samples transferred to a 37°C with 5% CO_2_ humid chamber.

#### In vitro microtubule nucleation assay

To investigate how MT nucleation was influenced by the presence of synaptic proteins, reaction chambers were prepared as described in (*64*). Synaptosomes were immobilised as described above. Atto-561-tubulin was used instead of HyLite tubulin at a concentration of 15 µM, and casein was substituted with 1% BSA for washes and blocking (***S***2C). An alexa488-labelled PSD95 was used to label synaptosomes. Images were recorded on a system described in Ramirez-Rios (*65*). For each condition, at least two slides from two independent experiments using different synaptosome preparations were done. Microtubule count was performed after an average of 30-40 min of incubation.

#### STED imaging

STED nanoscopy was performed using a Leica SP8-3xSTED and an Abberior Facility line imaging system. The Leica SP8-3 × STED microscope was used for the acquisition of three colour STED images. A white light laser was used for excitation at the wavelengths 488 nm for Alexa-488, 580 nm for Abberior-STAR-580, and 650 nm for Atto-647N. Fluorophore depletion was achieved with a 775 nm laser for AttoFluor-647N/Abberior-STAR-580 and a 592 nm laser for Alexa-488. All images were acquired using a 100×oil objective (Leica, HC APO CS2). Emission light was detected in bins: 660–730 nm for Atto-647N, 590–620 nm for AbberiorSTAR-580, and 500–530 nm for Alexa-488. Gated detection was applied with a delay of 0.3–1.5 ns. Pinhole size was set to 1 AU. Images were acquired with a 5 × zoom resulting in a pixel size of 22.73 nm, 1024 by 1024 and a 16 × line averaging.

GatedSTED images were acquired at a Leica TCS SP8-3X gatedSTED system (Leica Microsystems) equipped with a pulsed white light laser (WLL) for excitation ranging from 470 to 670 nm. STED was obtained with a 592 nm continuous wave and a 775 nm pulsed depletion laser. Samples were imaged with either a 100× objective (Leica, HC APO CS2 100×/1.40 oil) or a 93× glycerol objective (Leica, HC APO 93×/1.30 GLYC motCORR). The refractive index (RI) of Mowiol (see immunocytochemistry) polymerised for 3 days was 1.46 (measured with Digital Refractometer AR200 (Reichert), and matched the RI of glycerol (1.45) better than oil (1.51). Therefore, when available, the 93× glycerol objective was the preferred objective. For excitation of the respective channels, the WLL was set to 650 nm for phalloidin-Atto647N, 561 nm for Abberior STAR 580, and 488 nm for either Alexa Fluor 488-conjugated secondary antibodies or GFP-fusion proteins. STED was attained with the 775 nm laser for Atto647N/Abberior 580 and with the 592 nm laser for Alexa Fluor 488/eGFP. Emission spectra were detected at 660–730 nm for Atto647N, 580-620 nm for Abberior STAR 580, and 500–530 nm for Alexa Fluor 488 or eGFP. For gatedSTED, detector time gates were set to 0.5–6 ns for Abberior STAR 580/Atto647N and 1.5–6 ns for Alexa Fluor 488/eGFP. Images were acquired as single planes of either 1,024 × 1,024 pixels or 1,386 × 1,386, optical zoom factor 5 (for oil: x/y 22.73 nm, for glycerol: x/y 24.44 nm or 18.28 nm) or 6 (glycerol x/y 20.37 nm), 600 lines per second, and 16× line averaging. Corresponding confocal channels had the same setting as STED channels, except the excitation power was reduced and the detection time gates were set to 300 ps – 6 ns for all channels.

Additional STED imaging was performed on an Abberior STED Facility line microscope (Abberior Instruments GmbH) with an UPLSAPO 100 × oil immersion objective lens (NA 1.4). Pixel size was set to 20 nm for all images. Images were obtained from a 16 × frame accumulation. Excitation was achieved with pulsed diode lasers PDL-T 488, 561 and 640. Both the red and far-red channel were depleted using a 775 nm laser (PFL-40-3000-775-B1R). Pinhole size was set to 1AU. Gated detection was applied for both channels.

tauSTED microcopy was performed using a Leica STELLARIS 8 STED Falcon system (Leica) equipped with a WLL and a 660 and 775 nm depletion laser. Samples were imaged using a HC PL APO 93x/1.30 NA glycerol motCORR objective. HyD X detectors were used for FLIM and tauSTED imaging.

### Electrophysiological recordings

For electrophysiological measurements, we used 2-4 month old male SHANK3_L68P_ KI and SHANK3_WT_ littermates aged. The experimenter was blind to the genotypes. Animals were anesthetised with CO_2_ and decapitated. Brains were rapidly removed from the skull and placed in an ice-cold modified artificial cerebrospinal fluid solution (aCSF) containing (in mM): 110 choline chloride, 25 NaHCO_3_, 25 D-glucose, 11.6 Na-L-ascorbate, 7 MgSO_4_, 1.25 NaH_2_PO_4_, 2.5 KCl, 0.5 CaCl_2_ (pH 7.4 equilibrated with 95% O_2_ and 5% CO_2_). 400 µm thick transversal brain slices were prepared with a microtome vf-200 tissue slicer (Compresstome, Precisionary Instruments, USA) and then incubated at 30°C for about 90 min in physiologic aCSF containing (in mM): 124 NaCl, 26 NaHCO_3_, 10 D-glucose, 1 MgSO_4_, 1 NaH_2_PO_4_, 4 KCl, 2.4 CaCl_2_ (pH 7.4 equilibrated with 95% O_2_ and 5% CO_2_). After the incubation time, the hemi-slices were transferred to Synchroslice (Lohmann Research Equipment) recording chambers perfused with aCSF at a flow rate of ∼2 ml/min using a peristaltic pump (minipulse 3, Gilson, USA). The experiments were performed at 30°C. All recordings were executed by using a 2-channel Miniature Preamplifier (multichannel systems, Germany). The extracellular field excitatory post synaptic potentials (fEPSPs) recordings were performed by using a single fibre electrode (Lohmann Research Equipment, Germany) placed in the middle third of stratum radiatum in area CA1. fEPSPs were evoked by stimulating the Shaffer collateral (SC) fibres using a semi-micro concentric bipolar electrode (Lohmann Research Equipment, Germany) placed in the middle third of stratum radiatum 150-200 µm away from the recording electrode and approximately the same slice depth. Square-wave current pulses were generated by a stimulus generator (multichannel systems STG 4008, Germany) and delivered to the tissue. Input/Output function was generated by stimulating the SC fibers in 100 µA steps from 0 to 1500 µA.

Paired-Pulse ratio (PPR) was measured by delivering two stimuli for five times 30 seconds apart at 50, 100, 200, 500 ms inter-stimulus intervals. The PPR was calculated by dividing the amplitude of the second EPSP by the amplitude of the first EPSP. Synaptic facilitation was examined by repetitive stimulation (5 times) for each inter-stimulus interval, and the resulting potentials were averaged.

Recordings were analysed by using the SynchroSlice (Lohmann Research Equipment) software. Statistical comparisons of pooled data were performed by two-way ANOVA.

### Image analysis

Colocalisation of vGlut1 and SHANK3 was determined using ComDet v 0.4.2 in FIJI/ImageJ (Table 6). Maximum projections of both channels were used to detect puncta and ROIs were drawn around them. The overlap between both channels was measures in both directions (vGlut1/SHANK3 and SHANK3/vGlut1).

γ-tubulin positive PSD95 postsynapse in PV vs nonPV neurons were determined using ComDet v.0.5.5. The same dataset was used to determine the density of PSD95 postsynapses found on PV vs nonPV neurons. Similarly, SHANK3 puncta of SHANK3_L68P_ and SHANK3_WT_ dendrites were determined and the ROIs checked for tubulin intensity, determining synaptic microtubule content.

Colocalisation of CAMSAP2 with SHANK3 was analysed using OpenView (*66*). Puncta were detected in maximum projections of both channels and boxes drawn around their maxima using the Box_puncta_ex function. To determine colocalisation, the overlap between the two channels was measured using the “Match_Set1” function. Mean fluorescence intensities were determined in ROIs of both SHANK3 and CAMSAP2 channels. The threshold was then applied to all following images.

The percentage of PV/CB/CR neurons was analysed by manually counting MAP2+ cells, placing a ROI on their somata, yielding the number of neurons. The mean gray value of different GABAergic markers was determined in the ROIs. Cells positive for the individual markers were selected in the individual channels and the lowest mean gray value set as a threshold.

The density of F-actin patches in PV vs pCaMKII+ neurons was determined using the ImageJ/FIJI built-in function Analyze Particles.

The distance of the starting point of MacF18 tracks to PSD95 spots was measured using kymographs that were generated with the ImageJ plugin KymoClear by tracing the dendrite. Dynamic microtubule polarity was determined the same way.

In *in vitro* assays, colocalisation of synaptosomes with tubulin was determined manually. First, PSD95 puncta were indicated using the multi-point tool of ImageJ/FIJI. Then, it was confirmed whether the saved position was positive for tubulin.

To analyse the effect of the SHANK3_L68P_ mutation on dendritic branching of PV neurons, the dendritic arbour was traced using the ImageJ/FIJI plugin NeuronJ. An automated analysis using the ImageJ/FIJI plugin Simple Neurite Tracer (SNT) was performed. To display representative traces, ROIs were drawn onto a canvas.

To analyse the effect of CAMSAP2 MBD-expression on the dendritic morphology of PV and nonPV neurons, and to analyse the effect of CAMSAP2 depletion on the dendritic morphology of PV neurons expressing L68P-mutated or wildtype SHANK3, the dendritic arbour was traced using SNT.

Immunoblots were analysed using ImageJ/FIJI-built in function GelAnalyzer or if at least one sample did not contain a band by measuring mean gray values of same-size ROIs.

### Statistics

When relying on manual analyses, the person analysing the data was generally blinded, if possible. Microscopy and immunoblotting images are displayed with identical settings when obtained from the same preparation when comparing e.g. intensities between groups.

Graphpad prism was used for statistical analyses (Table 6). Data are presented as mean ± SEM. Before parametric statistical analysis, we used the one-sample Kolmogorov-Smirnov and D’Agostino & Pearson test to assess the normal distribution of our datasets. When comparing two groups, two-tailed t- or Mann-Whitney U tests were used when data was or was not normal distributed, respectively. Graphpad Prism was used for all statistical analyses.

## Acknowledgments

This research was supported by: German research foundation (Excellence Strategy – EXC-2049–390688087 to MM, DFG CRC1315 project A03 to MM, DFG FOR5228 to MM, DFG FOR2419 to MM, the Institute for Basic Science IBS-R002-D1 to EK. Leducq Foundation, research grant n° 20CVD01 to MJM. Mass spectrometry data were generated using an instrument funded through the DFG (project number 428987924 to K.K.). We would further like to thank the Interdisciplinary Center Life in Time and Space (IZ LIST) for awarding a travel grant to support a stay at EMBL Imaging Centre to DH. We would like to thank the EMBL Imaging Centre and especially Timo Zimmermann and Dietrich Walsh for access to the STELLARIS 8 STED Falcon and the STELLARIS STED microscope for DH, and the Photonic Imaging Center of Grenoble Institute Neuroscience (Univ Grenoble Alpes – Inserm U1216), which is part of the ISdV core facility. For providing CAMSAP2 plasmids, we would like to thank Anna Akhmanova and Kai Jiang. We would like to thank Thao Thi Bich Nguyen, Lisa Mallis and Minji Kim for technical support.

## Author contributions

Conceptualisation: DH, MM

Methodology: DH, MHB, TF, TY, SRR

Investigation: DH, SuL, MHB, SaL, NAL, TF, CL, AMB, SRR, CS, DHWD, MM

Visualisation: DH, CL

Supervision: KK, JAAD, MJM, EK, MM

Writing - original draft: DH

Writing - review & editing: SuL, MHB, TF, AMB, SRR, TY, MJM, EK, MM

## Competing interests

The authors declare no conflict of interest.

## Data and materials availability

Data are available in the main text or the supplementary materials. Mass spectrometry results will be uploaded to the PRIDE databank.

## References

1. M. M. Rolls, P. Thyagarajan, C. Feng, Microtubule dynamics in healthy and injured neurons. Dev Neurobiol., 13 (2021).

2. A. Sakakibara, R. Ando, T. Sapir, T. Tanaka, Microtubule dynamics in neuronal morphogenesis. Open Biol. 3 (2013).

3. F. Borys, E. Joachimiak, H. Krawczyk, H. Fabczak, Molecules25163705. 1–36 (2020).

4. A. J. Baines, P. A. Bignone, M. D. A. King, A. M. Maggs, P. M. Bennett, J. C. Pinder, G. W. Phillips, The CKK domain (DUF1781) binds microtubules and defines the CAMSAP/ssp4 family of animal proteins. Mol. Biol. Evol. 26, 2005–2014 (2009).

5. K. J. Verhey, J. Gaertig, The tubulin code. Cell Cycle 6, 2152–2160 (2007).

6. R. P. Tas, A. Chazeau, B. M. C. Cloin, M. L. A. Lambers, C. C. Hoogenraad, L. C. Kapitein, Differentiation between Oppositely Oriented Microtubules Controls Polarized Neuronal Transport. Neuron 96, 1264–1271.e5 (2017).

7. M. K. Iwanski, L. C. Kapitein, Cellular cartography: Towards an atlas of the neuronal microtubule cytoskeleton. Front. Cell Dev. Biol. 11 (2023).

8. S. F. B. Van Beuningen, L. Will, M. Harterink, A. Chazeau, E. Y. Van Battum, C. P. Frias, M. A. M. Franker, E. A. Katrukha, R. Stucchi, K. Vocking, A. T. Antunes, L. Slenders, S. Doulkeridou, P. Sillevis Smitt, A. F. M. Altelaar, J. A. Post, A. Akhmanova, R. J. Pasterkamp, L. C. Kapitein, E. de Graaff, C. C. Hoogenraad, TRIM46 Controls Neuronal Polarity and Axon Specification by Driving the Formation of Parallel Microtubule Arrays. Neuron 88, 1208–1226 (2015).

9. D. J. Sharp, W. Yu, L. Ferhat, R. Kuriyama, D. C. Rueger, P. W. Baas, Identification of a microtubule-associated motor protein essential for dendritic differentiation. Journal of Cell Biology 138, 833–843 (1997).

10. S. Lin, M. Liu, O. I. Mozgova, W. Yu, P. W. Baas, Mitotic motors coregulate microtubule patterns in axons and dendrites. Journal of Neuroscience 32, 14033–14049 (2012).

11. M. Schelski, F. Bradke, Microtubule retrograde flow retains neuronal polarization in a fluctuating state. Sci. Adv. 8, 1–19 (2022).

12. K. W. Yau, P. Schätzle, E. Tortosa, S. Pagès, A. Holtmaat, L. C. Kapitein, C. C. Hoogenraad, Dendrites In vitro and In vivo contain microtubules of opposite polarity and axon formation correlates with uniform plus-end-out microtubule orientation. Journal of Neuroscience 36, 1071–1085 (2016).

13. P. W. Baas, J. S. Deitch, M. M. Black, G. A. Banker, Polarity orientation of microtubules in hippocampal neurons: Uniformity in the axon and nonuniformity in the dendrite. Proc. Natl. Acad. Sci. U. S. A. 85, 8335–8339 (1988).

14. A. Mukherjee, P. S. Brooks, F. Bernard, A. Guichet, P. T. Conduit, Microtubules originate asymmetrically at the somatic golgi and are guided via kinesin2 to maintain polarity within neurons. Elife 9, 1–33 (2020).

15. A. Yagoubat, P. T. Conduit, Asymmetric microtubule nucleation from Golgi stacks promotes opposite microtubule polarity in axons and dendrites. Current Biology 35, 1311–1325.e4 (2025).

16. A. T. Weiner, P. Thyagarajan, Y. Shen, M. M. Rolls, To nucleate or not, that is the question in neurons. Neurosci. Lett. 751, 135806 (2021).

17. M. Stiess, N. Maghelli, L. C. Kapitein, S. Gomis-Rüth, M. Wilsch-Bräuninger, C. C. Hoogenraad, I. M. Tolić-Nørrelykke, F. Bradke, Axon extension occurs independently of centrosomal microtubule nucleation. Science (1979). 327, 704–707 (2010).

18. K. M. Ori-McKenney, L. Y. Jan, Y. N. Jan, Golgi Outposts Shape Dendrite Morphology by Functioning as Sites of Acentrosomal Microtubule Nucleation in Neurons. Neuron 76, 921–930 (2012).

19. A. T. Weiner, D. Y. Seebold, P. Torres-Gutierrez, C. Folker, R. D. Swope, G. O. Kothe, J. G. Stoltz, M. K. Zalenski, C. Kozlowski, D. J. Barbera, M. A. Patel, P. Thyagarajan, M. Shorey, D. M. R. Nye, M. Keegan, K. Behari, S. Song, J. D. Axelrod, M. M. Rolls, Endosomal Wnt Signaling Proteins Control Microtubule Nucleation in Dendrites (2020; 10.1371/journal.pbio.3000647)vol. 18.

20. M. C. Stone, F. Roegiers, M. M. Rolls, Microtubules have opposite orientation in axons and dendrites of Drosophila neurons. Mol. Biol. Cell 19, 4122–4129 (2008).

21. N. Nakagawa, The neuronal Golgi in neural circuit formation and reorganization. Front. Neural Circuits 18 (2024).

22. C. Sanyal, N. Pietsch, S. Ramirez Rios, L. Peris, L. Carrier, M. J. Moutin, The detyrosination/re-tyrosination cycle of tubulin and its role and dysfunction in neurons and cardiomyocytes. Semin. Cell Dev. Biol. 137, 46–62 (2023).

23. J. Gu, B. L. Firestein, J. Q. Zheng, Microtubules in dendritic spine development. Journal of Neuroscience 28, 12120–12124 (2008).

24. L. C. Kapitein, K. W. Yau, C. C. Hoogenraad, “Chapter 7 - Microtubule Dynamics in Dendritic Spines” in Methods in Cell Biology, L. Cassimeris, P. Tran, Eds. (Academic Press, 2010; https://www.sciencedirect.com/science/article/pii/S0091679X10970076)vol. 97 of *Microtubules: in vivo*, pp. 111–132.

25. K. Wai Yau, S. F. van Beuningen, I. Cunha-Ferreira, B. M. Cloin, E. Y. van Battum, L. Will, P. Schä tzle, R. P. Tas, J. van Krugten, E. A. Katrukha, K. Jiang, P. S. Wulf, M. Mikhaylova, M. Harterink, R. Jeroen Pasterkamp, A. Akhmanova, L. C. Kapitein, C. C. Hoogenraad, Article Microtubule Minus-End Binding Protein CAMSAP2 Controls Axon Specification and Dendrite Development. Neuron 82, 1058–1073 (2014).

26. M. Megías, Z. Emri, T. F. Freund, A. I. Gulyás, Total number and distribution of inhibitory and excitatory synapses on hippocampal CA1 pyramidal cells. Neuroscience 102, 527–540 (2001).

27. L. Nahar, B. M. Delacroix, H. W. Nam, The Role of Parvalbumin Interneurons in Neurotransmitter Balance and Neurological Disease. Front. Psychiatry 12, 679960 (2021).

28. Y. S. Hwang, C. Maclachlan, J. Blanc, A. Dubois, C. C. H. Petersen, G. Knott, S. H. Lee, 3D Ultrastructure of Synaptic Inputs to Distinct GABAergic Neurons in the Mouse Primary Visual Cortex. Cerebral Cortex (New York, NY) 31, 2610 (2021).

29. C. A. Runyan, M. Sur, Response selectivity is correlated to dendritic structure in parvalbumin-expressing inhibitory neurons in visual cortex. Journal of Neuroscience 33, 11724–11733 (2013).

30. W. Feng, M. Zhang, Organization and dynamics of PDZ-domain-related supramodules in the postsynaptic density. Nature Publishing Group [Preprint] (2009). 10.1038/nrn2540.

31. M. Sheng, C. C. Hoogenraad, The Postsynaptic Architecture of Excitatory Synapses: A More Quantitative View. Annu. Rev. Biochem. 76, 823–847 (2007).

32. H.-J. Kreienkamp, “Scaffolding Proteins at the Postsynaptic Density: Shank as the Architectural Framework” in Protein-Protein Interactions as New Drug Targets, E. Klussmann, J. Scott, Eds. (Springer, Berlin, Heidelberg, 2008; 10.1007/978-3-540-72843-6_15)*Handbook of Experimental Pharmacology*, pp. 365–380.

33. M. Bucher, T. Fanutza, M. Mikhaylova, Cytoskeletal makeup of the synapse: Shaft versus spine. Cytoskeleton 77, 55–64 (2020).

34. C. M. Durand, C. Betancur, T. M. Boeckers, J. Bockmann, P. Chaste, F. Fauchereau, G. Nygren, M. Rastam, I. C. Gillberg, H. Anckarsäter, E. Sponheim, H. Goubran-Botros, R. Delorme, N. Chabane, M. C. Mouren-Simeoni, P. De Mas, E. Bieth, B. Rogé, D. Héron, L. Burglen, C. Gillberg, M. Leboyer, T. Bourgeron, Mutations in the gene encoding the synaptic scaffolding protein SHANK3 are associated with autism spectrum disorders. Nat. Genet. 39, 25–27 (2007).

35. L. Boccuto, M. Lauri, S. M. Sarasua, C. D. Skinner, D. Buccella, A. Dwivedi, D. Orteschi, J. S. Collins, M. Zollino, P. Visconti, B. Dupont, D. Tiziano, R. J. Schroer, G. Neri, R. E. Stevenson, F. Gurrieri, C. E. Schwartz, Prevalence of SHANK3 variants in patients with different subtypes of autism spectrum disorders. European Journal of Human Genetics 21, 310–316 (2013).

36. J. Gauthier, D. Spiegelman, A. Piton, R. G. Lafrenière, S. Laurent, J. St-Onge, L. Lapointe, F. F. Hamdan, P. Cossette, L. Mottron, É. Fombonne, R. Joober, C. Marineau, P. Drapeau, G. A. Rouleau, Novel de novo SHANK3 mutation in autistic patients. American Journal of Medical Genetics Part B: Neuropsychiatric Genetics 150B, 421–424 (2009).

37. C. M. Durand, J. Perroy, F. Loll, D. Perrais, L. Fagni, T. Bourgeron, M. Montcouquiol, N. Sans, SHANK3 mutations identified in autism lead to modification of dendritic spine morphology via an actin-dependent mechanism. Mol. Psychiatry 17, 71–84 (2012).

38. Y. E. Yoo, T. Yoo, S. Lee, J. Lee, D. Kim, H. M. Han, Y. C. Bae, E. Kim, Shank3 mice carrying the human q321r mutation display enhanced self-grooming, abnormal electroencephalogram patterns, and suppressed neuronal excitability and seizure susceptibility. Front. Mol. Neurosci. 12, 1–23 (2019).

39. M. Bucher, S. Niebling, Y. Han, D. Molodenskiy, F. H. Nia, H. J. Kreienkamp, D. Svergun, E. Kim, A. S. Kostyukova, M. R. Kreutz, M. Mikhaylova, Autism-associated SHANK3 missense point mutations impact conformational fluctuations and protein turnover at synapses. Elife 10 (2021).

40. R. Palenzuela, Y. Gutiérrez, J. E. Draffin, A. Lario, M. Benoist, J. A. Esteban, MAP1B light chain modulates synaptic transmission via AMPA receptor intracellular trapping. Journal of Neuroscience 37, 9945–9963 (2017).

41. J. Parato, F. Bartolini, The microtubule cytoskeleton at the synapse. Neurosci. Lett. 753 (2021).

42. O. Yagensky, T. K. Dehaghi, J. J. E. Chua, The roles of microtubule-based transport at presynaptic nerve terminals. Front. Synaptic Neurosci. 8, 1–9 (2016).

43. E. W. Dent, Dynamic microtubules at the synapse. Elsevier Ltd [Preprint] (2020). 10.1016/j.conb.2020.01.003.

44. L. C. Kapitein, K. W. Yau, C. C. Hoogenraad, Microtubule Dynamics in Dendritic Spines. Methods Cell Biol. 97, 111–132 (2010).

45. R. P. Tas, L. C. Kapitein, Exploring cytoskeletal diversity in neurons. Science (1979). 361, 231–232 (2018).

46. Y. Aika, J. Q. Ren, K. Kosaka, T. Kosaka, Quantitative analysis of GABA-like-immunoreactive and parvalbumin-containing neurons in the CA1 region of the rat hippocampus using a stereological method, the disector. Exp. Brain Res. 99, 267–276 (1994).

47. W. Woodson, L. Nitecka, Y. Ben-Ari, Organization of the GABAergic system in the rat hippocampal formation: a quantitative immunocytochemical study. J. Comp. Neurol. 280, 254–271 (1989).

48. D. Keller, C. Erö, H. Markram, Cell densities in the mouse brain: A systematic review. Front. Neuroanat. 12, 394837 (2018).

49. Y. Han, D. Hacker, B. C. Donders, C. Parperis, R. Thuenauer, C. Leterrier, K. Grünewald, M. Mikhaylova, Unveiling the cell biology of hippocampal neurons with dendritic axon origin. J Cell Biol J Cell Bio (2025).

50. M. M. Rolls, Principles of microtubule polarity in linear cells. Dev. Biol. 483, 112–117 (2022).

51. V. Sulimenko, E. Dráberová, P. Dráber, γ-Tubulin in microtubule nucleation and beyond. Front. Cell Dev. Biol. 10, 1–14 (2022).

52. M. M. Nguyen, C. J. McCracken, E. S. Milner, D. J. Goetschius, A. T. Weiner, M. K. Long, N. L. Michael, S. Munro, M. M. Rolls, Gamma-tubulin controls neuronal microtubule polarity independently of Golgi outposts. Mol Biol Cell 25, 2039–2050 (2014).

53. J. P. Vessey, D. Karra, More than just synaptic building blocks: Scaffolding proteins of the post-synaptic density regulate dendritic patterning. J. Neurochem. 102, 324–332 (2007).

54. T. M. Böckers, M. G. Mameza, M. R. Kreutz, J. Bockmann, C. Weise, F. Buck, D. Richter, E. D. Gundelfinger, H. J. Kreienkamp, Synaptic scaffolding proteins in rat brain: Ankyrin repeats of the multidomain Shank protein family interact with the cytoskeletal protein α-fodrin. Journal of Biological Chemistry 276, 40104–40112 (2001).

55. B. van Bommel, A. Konietzny, O. Kobler, J. Bär, M. Mikhaylova, F-actin patches associated with glutamatergic synapses control positioning of dendritic lysosomes. EMBO J. 38 (2019).

56. P. Thyagarajan, C. Feng, D. Lee, M. Shorey, M. M. Rolls, Microtubule polarity is instructive for many aspects of neuronal polarity. Dev. Biol. 486, 56–70 (2022).

57. Michael Bucher, Characterization of the Structural and Functional Impact of Autism-Associated SHANK3 Missense Mutations (2022).

58. K. Jiang, S. Hua, R. Mohan, I. Grigoriev, K. W. Yau, Q. Liu, E. A. Katrukha, A. F. M. Altelaar, A. J. R. Heck, C. C. Hoogenraad, A. Akhmanova, Microtubule Minus-End Stabilization by Polymerization-Driven CAMSAP Deposition. Dev. Cell 28, 295–309 (2014).

59. Q. Chang, H. Yang, M. Wang, H. Wei, F. Hu, Role of Microtubule-Associated Protein in Autism Spectrum Disorder. Neurosci. Bull. 34, 1119–1126 (2018).

60. C. S. Leblond, C. Nava, A. Polge, J. Gauthier, G. Huguet, S. Lumbroso, F. Giuliano, C. Stordeur, C. Depienne, K. Mouzat, D. Pinto, J. Howe, N. Lemière, C. M. Durand, J. Guibert, E. Ey, R. Toro, H. Peyre, A. Mathieu, F. Amsellem, M. Rastam, I. C. Gillberg, G. A. Rappold, R. Holt, A. P. Monaco, E. Maestrini, P. Galan, D. Heron, A. Jacquette, A. Afenjar, A. Rastetter, A. Brice, F. Devillard, B. Assouline, F. Laffargue, J. Lespinasse, J. Chiesa, F. Rivier, D. Bonneau, B. Regnault, D. Zelenika, M. Delepine, M. Lathrop, D. Sanlaville, C. Schluth-Bolard, P. Edery, L. Perrin, A. C. Tabet, M. J. Schmeisser, T. M. Boeckers, M. Coleman, D. Sato, P. Szatmari, S. W. Scherer, G. A. Rouleau, C. Betancur, M. Leboyer, C. Gillberg, R. Delorme, T. Bourgeron, Meta-analysis of SHANK Mutations in Autism Spectrum Disorders: A Gradient of Severity in Cognitive Impairments. PLoS Genet. 10 (2014).

61. T. Yoo, H. Cho, J. Lee, H. Park, Y. E. Yoo, E. Yang, J. Y. Kim, H. Kim, E. Kim, GABA neuronal deletion of Shank3 exons 14–16 in mice suppresses striatal excitatory synaptic input and induces social and locomotor abnormalities. Front. Cell. Neurosci. 12, 1–16 (2018).

62. J. Pagano, S. Landi, A. Stefanoni, G. Nardi, M. Albanesi, H. F. Bauer, E. Pracucci, M. Schön, G. M. Ratto, T. M. Boeckers, C. Sala, C. Verpelli, Shank3 deletion in PV neurons is associated with abnormal behaviors and neuronal functions that are rescued by increasing GABAergic signaling. Mol. Autism 14, 28 (2023).

63. M. Mikhaylova, J. Bär, B. van Bommel, P. Schätzle, P. A. YuanXiang, R. Raman, J. Hradsky, A. Konietzny, E. Y. Loktionov, P. P. Reddy, J. Lopez-Rojas, C. Spilker, O. Kobler, S. A. Raza, O. Stork, C. C. Hoogenraad, M. R. Kreutz, Caldendrin Directly Couples Postsynaptic Calcium Signals to Actin Remodeling in Dendritic Spines. Neuron, doi: 10.1016/j.neuron.2018.01.046 (2018).

64. V. Stoppin-Mellet, N. Bagdadi, Y. Saoudi, I. Arnal, “Studying Tau-Microtubule Interaction Using Single-Molecule TIRF Microscopy” in Cytoskeleton Dynamics (2019), pp. 77–91.

65. S. Ramirez-Rios, S. R. Choi, C. Sanyal, T. B. Blum, C. Bosc, F. Krichen, E. Denarier, J. M. Soleilhac, B. Blot, C. Janke, V. Stoppin-Mellet, M. M. Magiera, I. Arnal, M. O. Steinmetz, M. J. Moutin, VASH1–SVBP and VASH2–SVBP generate different detyrosination profiles on microtubules. Journal of Cell Biology 222 (2023).

66. S. Tsuriel, R. Geva, P. Zamorano, T. Dresbach, T. Boeckers, E. D. Gundelfinger, C. C. Garner, N. E. Ziv, Local sharing as a predominant determinant of synaptic matrix molecular dynamics. PLoS Biol. 4, 1572–1587 (2006).

